# Damage-related changes in the cerebellum of juvenile *Oncorhynchus masou*: reactivation of neurogenic niches and astrocytic response

**DOI:** 10.1101/681445

**Authors:** Eugenyia V. Pushchina, Maryia E. Stukaneva, Anatoly A. Varaksin

**Author notes:** Abbreviation: CCb, *corpus cerebelli*; DMZ, dorsal matrix zone; GS, glutamine synthetase; PC, Purkinje cells; EDC, eurydendroid cells.

## Abstract

In the cerebellum of juvenile *Oncorhynchus masou*, proliferating BrdU+ and HuCD+ cells and constitutive neurogenic niches were detected in different zones; the largest number of labeled cells were found in the dorsal part of the molecular layer and the dorsal matrix zone (DMZ). Cells labeled with glutamine synthetase (GS) and radial glia were also present in the intact *O. masou* cerebellum. The most intensive proliferation was detected in the rostral part of cerebellum. This part is assumed to contain active zones of constitutive neurogenesis. After an injury inflicted to the cerebellum, the number of BrdU+ and HuCD+ cells increased significantly. The number of BrdU+ cells after this type of injury was much greater than after a telencephalon trauma. A quantitative analysis revealed that after the cerebellum injury the proliferative activity in the caudal part of CCb is increased compared to that in the control. A reactivation of neurogenic and neuroepithelial niches and their transformation into reactive neurogenic domains, with an increased distribution density of intensely labeled HuCD+ cells of different types, were observed. The increase in the number of HuCD+ differentiated cells in the basal area suggests that the processes of neuronal differentiation are intensified in the cerebellum of juvenile *O. masou* after injury. The number of GS positive cells (GS+) and fibers increased in all the zones of cerebellum. The most intensive astrocytic response was noted in the dorsal part of cerebellum. The data of the enzyme immunoassay confirm the multiple variations in the level of GS after a traumatic injury to cerebellum in *O. masou*.

## 1. Introduction

Fish brain, growing with the body throughout the life, is an interesting model for studying the brain recovery after a traumatic injury and the influence of various factors on these processes. In an adult mammalian brain, acute inflammation interferes with adult neurogenesis and regeneration (Hoehn et al., 2005; Iosif et al., 2006), but in the brain of an adult zebrafish, *Danio rerio*, the inflammation process can contribute to the recovery of the central nervous system (CNS) after trauma (Kyritsis et al., 2012). A recent study on adult *D. rerio* has shown that the inflammatory response is necessary for the initiation of neurogenic proliferation in the ventricular zone, since it initiates a number of molecular signals preceding the activation of radial glia necessary to enhance proliferation and reparative neurogenesis (Kyritsis et al., 2012). Various types of damage to fish brain create special conditions for the implementation of genetic programs that enhance the proliferation of progenitor cells, and also activate the neurogenic niches and the proliferation of neural stem cells (NSC) in them (Kishmoto et al., 2012; Kaslin et al., 2013).

Glutamate is an exciting neurotransmitter that, when interacting with membrane receptors, regulates numerous functions in the CNS, such as cognition, memory, movement, etc. A high extracellular level of glutamate, accumulated as a result of neuronal damage, causes many neurodegenerative diseases (Benarroch, 2010). Glutamine synthetase (GS) is a specific glial protein that converts neurotoxic glutamate into amino acid glutamine. This mechanism protects neurons from cell death (Struzyńska, 2009; Benarroch, 2010). In recent studies, it has been shown that GS labels radial glia in fish brain (Grupp et al., 2010). A brain trauma, focal ischemia, and a number of neurological diseases are often accompanied by a decrease in the expression of GS, enhancing the neurodegenerative effect caused by the glutamate neurotoxicity (Grosche et al., 1995). In the light of the finding of the reduction in GS after a damage to mammalian CNS, a number of studies (Struzyńska, 2009; Benarroch, 2010) provided data on a significant increase in GS in fish cerebellum after a mechanical injury, which are of particular interest (Zupanc and Sîrbulescu, 2013). GS converts toxic glutamate into neutral glutamine and plays an important role in the nitrogen metabolism in fish (Dhanasiri et al., 2012). It is possible that an increase in GS in fish provides an important mechanism for reducing neurodegenerative effects caused by the glutamate neurotoxicity (Zupanc, 2007).

In different parts of brain in salmonids, a high level of proliferation of cells was found in the ventricular zone (Pushchina et al., 2012, 2013). A significant number of proliferating cell nuclear antigen PCNA+ and HuCD+ cells were observed in the pallial periventricular zone of telencephalon in *O. masou* juveniles (Pushchina et al., 2017a). PCNA+ cells were found in the periventricular zone of diencephalon, mesencephalic tegmentum, and rhomboencephalon (Pushchina et al., 2012). Studies based on labeling of PCNA showed a high intensity of cell proliferation in the dorsal matrix zone (DMZ) of the cerebellum in juvenile *O. masou* (Stukaneva et al., 2017; Pushchina et al., 2018). This zone is located in the dorsomedial part of the cerebellum’s body, at the boundary between the molecular and granular layers.

In the dorsal, lateral, and basal zones of the molecular layer of cerebellum in *O. masou*, neurogenic niches containing PCNA+ cells and a heterogeneous population of PCNA cells were identified after injury (Stukaneva et al., 2017). The appearance of reactive neurogenic niches after a damaging impact is a feature of reparative neurogenesis in fish brain (Than-Trong and Bally-Cuif, 2015). A study of damage to the telencephalon of adult *D. rerio* has shown that persistent inflammation does not spread widely, and the glial scar does not develop (Baumgart et al., 2012). However, in the cerebellum of *O. masou* juveniles, radial glial fibers and single small cells, intensely and moderately labeled with glial fibrillary acidic protein (GFAP), were found after damage (Stukaneva et al., 2017). As a result of the damaging impact, GFAP+ radial glia fibers appeared in the dorsal matrix zone of cerebellum, forming multidirectional, radially oriented beams (Pushchina et al., 2018). No similar structural formations were detected in intact *O. masou* juveniles. After the cerebellum injury in juvenile *O. masou*, the dorsal matrix zone undergoes a structural rearrangement associated with a partial spatial reorientation of the radial glia fibers and the formation of specific guides for the cells formed in this zone (Stukaneva et al., 2017).

It has been shown that in zebrafish the proliferative activity and the response from the NSC can vary between different areas of the brain after damage (Grandel and Brandt, 2013). It is also now evident that the method of traumatic damage to the fish brain influences its reparative capacity (März et al., 2011; Baumgart et al., 2012). To further characterize the processes of proliferation in the cerebellum of juvenile *O. masou*, we used different variants of brain damage: mechanical injury to the cerebellum and to the telencephalon. In both variants, the inclusion of BrdU in cells of the damaged cerebellum was examined on day 3 post-injury. We studied the features of constitutive and reparative neurogenesis in the cerebellum of juvenile *O. masou*; the changes in the synthesis of GS after a mechanical injury were also investigated.

## 2. Material and methods

### 2.1. Animals and housing conditions

In the work, we used 70 *Oncorhynchus masou* juveniles with a body length of 9–11.5 cm and a weight of 20–35 g. The animals were obtained from the Ryazanovka experimental production fish hatchery (Primorsky Krai, Russia) in 2018D. The animals were kept in a 6 tanks with aerated fresh water at a temperature of 16–17°C and fed once a day. The size of the tanks was 90×60×50 cm. The light/dark cycle was 14/10 h. The dissolved oxygen content of water was 7–10 mg/dm^3^, which corresponds to normal saturation. All the experiments on animals were supervised and approved by the Commission on Biomedical Ethics of the National Scientific Center of Marine Biology (NSCMB), Far Eastern Branch, Russian Academy of Sciences (FEB RAS) (approval №. 4-031018).

### 2.2. Tissue Sampling

The fish were anesthetized in a 2% solution of tricaine methanesulfonate, MS222 (Sigma-Aldrich, USA), for 10–15 min. After anesthesia, the intracranial cavity of the immobilized animal was perfused with a 4% paraformaldehyde solution prepared in 0.1 M phosphate buffer (pH 7.2) using a syringe. After prefixation, the brain was removed from the skull cavity and fixed in a 4% paraformaldehyde solution for two hours at 4°C. Then it was kept in a 30% solution of sucrose at 4°C for two days (with five replacements of the solution). A series of frontal 50-μm-thick sections of the fish brain were made on a freezing microtome Cryo-star HM 560 MV (Carl Zeiss, Oberkochen, Germany), mounted on gelatinized slides, and dried. For the immunohistochemistry (IHC) of BrdU, GS, and HuCD, every third frontal section of the cerebellum was taken for examination. For morphological analysis, all the frontal sections of the *O. masou* cerebellum were stained with 1% aqueous solution of toluidine blue.

### 2.3. Mechanical damage to cerebellum and telencephalon. BrdU administration

The experimental damage to the cerebellum of *O. masou* juveniles was carried out according to the methods of Clint and Zupanc (2001). All chemicals were purchased from Sigma (St Louis, LO, USA). A mechanical injury was applied to the dorsalmost part of *corpus cerebelli* (CCb), as described by Clint and Zupanc (2001). The damage zone included both the dorsal molecular and the granular layers of the cerebellum’s body and did not affect other parts of the brain. Immediately after inflicting the injury, the animals were put into a tank with fresh water for recovery and further monitoring. Within the first hour post-injury, the animals showed increased locomotor activity (jumped out of the water and made rotary motions). These changes in motor activity of fish indicate an injury of the cerebellum structure.

The individuals were subjected to general anesthesia by immersion into a tank with 2% MS 222. A 3-mm deep stab wound, puncturing the cranium of fish, was inflicted in the parasagittal direction to CCb. In the other case, the injury was applied in a similar way to the right hemisphere of telencephalon by the method of Kishimoto with co-authors (Kishimoto et al., 2012). In both experimental groups (n = 5 for each group), concurrently with the cerebellum/telencephalon damage, the animals were administered an intraperitoneal injection of 10 mg/mL BrdU solution (Sigma-Aldrich, USA) at a dose of 20 µL/g body weight. After a post-injection survival period of 3 days, the fish were perfused intracardially, and their brain was removed for further preparation. Animals in the control group (n = 5) received only BrdU injection. Immediately after the mechanical damage, the animals were released into a сommon tank with fresh water for their recovery and further monitoring.

For IHC of GS and HuCD, the animals were divided into two groups (n = 5 for each group). The first group consisted of intact animals; the second group, the animals with the cerebellum injury, which were examined on day 3 post-injury.

### 2.4. Immunohistochemistry

The tissue sections were processed for BrdU immunohistochemistry (IHC). For anti-5-bromo-2’-deoxyuridine (BrdU) immunolabeling, 7-μm-thick paraffin embedded sections were de-embedded according to the standard histological protocol. Preparation of sections for IHC was carried out in accordance with the BrdU-staining protocol of Dolbeare (1995). The sections of brain were incubated in 1 M HCl for 10 min on ice to break open the DNA structure of the labeled cells. This was followed by incubation in 2 M HCl for 10 min at room temperature, then for 20 min at 37°C. Immediately after the acid incubations, the sections were neutralized by incubating in 0.1 M borate buffer for 10 min at room temperature and washed in PBS (pH 7.4), 0.1% Triton X-100, three times, 5 min per wash. A 1% hydrogen peroxide solution on 0.1 M phosphate buffer was applied to the sections (pH 7.2). The sections were then incubated for 20 min at room temperature and washed thrice in 0.1 M phosphate buffer for 5 minutes per wash. Subsequently, the sections were incubated with the anti-bromodeoxyuridine/BrdU monoclonal mouse antibody (1:200; clone SPM166; Novus Biologicals, Littleton, USA) at room temperature for 30 min and then washed in three shifts of 0.1 M phosphate buffer for 5 minutes per shift. To visualize IHC labeling, a standard Vectastain Elite ABC kit (Vector Laboratories, United States) was used according to the manufacturer’s instructions. The red substrate (VIP Substrate Kit, Vector Labs, Burlingame, United States) was used for visualization of IHC reaction. The sections were dehydrated according to a standard procedure and enclosed under coverslips in the BioOptica Milano Spa (Italy) medium for concluding mounts.

To identify HuCD and GS, frontal 50-μm brain sections were cut on a Cryo-star HM 560 MV (Carl Zeiss, Oberkochen, Germany) freezing microtome. The brain sections were dehydrated for 90 min and washed thrice in 0.1 M phosphate-buffered saline. Immunohistochemical revelation of HuCD and GS was performed using standard avidin-biotin peroxidase labeling on freely floating sections. The sections of cerebellum were incubated with monoclonal mouse antibodies against HuCD (ICN, Biomedicals, United States) (1:300) and glutamine synthetase (Vector Laboratories Burlingame, United States) (1:300) at 4°C for 48 h. To visualize the immunohistochemical labeling, a standard kit (Vectastain Elite ABC Kit, Burlingame, United States) was used. The red substrate (VIP Substrate Kit, Vector Labs, Burlingame, United States) in combination with methyl green staining according to Brachet’s technique was applied. The development of the IHC staining was monitored under a microscope. The sections were dehydrated by a standard protocol and embedded in the medium (Bio-Optica, Italy). The mounts were examined and photographed on the Axiovert Apotome microscope.

The specificity of the immunohistochemical reaction was evaluated using a method of negative control. The brain sections were incubated with 1% non-immune horse serum, instead of primary antibodies, for 24 h and then studied as those with primary antibodies. In all the control experiments, the immunopositive reaction was absent.

### 2.5. ELISA immunoassay of the level of glutamine synthetase in O. masou cerebellum after injury

ELISA immunoassay of the quantitative level of GS in the brain of intact fish and fish after the cerebellum injury was performed using a Fish Glutamine Synthetase ELISA Kit (MBS011386, USA). For the enzyme immunoassay, we used the material from intact animals and fish at 1h, 3h, 12h, 1d, 2d, 3d, and 5 days after the mechanical injury of cerebellum (n = 3 in each group).

The brain of intact and experimental fish was removed from the skull in a 0.02M phosphate buffer, weighed and then thoroughly washed in an ice-cooled 0.02 M phosphate buffer (pH 7.2) to remove blood. The brain tissue was further mechanically cut into small pieces in 5 mL of phosphate buffer in a Potter-Elvehim PTFE glass homogenizer (Sigma, Aldrich, USA) on ice. The *O. masou* brain homogenates contained 10 mg tissue per 100 μL of PBS. The resulting suspension was sonicated on a Sonoplus 2070 ultrasonic homogenizer (Bandelin, Germany, Berline) to destroy cell membranes. The homogenates were then centrifuged for 15 minutes at 1500 × g (5000 rpm) in a rotor (Beckman Coulter Ti50, USA, CA, Palo Alto). The supernatant was analyzed using a Fish Glutamine Synthetase ELISA Kit (MBS, USA) immunoperoxidase identification system according to the manufacturer’s protocol. A standard solution was used for standardization. The assay was carried out in a proprietary 96-well microtiter plate. Optical density was measured at a wavelength of 450 nm for 15 min with a densitometer (Microelisa Stripplate Reader, Bio-Rad, USA).

### 2.6. Morphometric analysis

Morphometric analysis was performed using the software of an Axiovert 200 M inverted microscope equipped with the ApoTome module and Axio Cam MRM and Axio Cam HRC (Carl Zeiss, Germany) digital cameras. Microphotographs of sections were processed and the sample analysis was conducted using the AxioVision software. Measurements were performed at a 400× magnification in five randomly chosen fields of view for each area examined and an average value was calculated.

### 2.7. Statistical analysis

The proliferative index was determined by counting the number of positively stained cells among 100 nuclei (percentage) from five randomly selected view fields at 200× magnification, using the Axiovision software (Carl Zeiss, Germany).

Quantitative processing of the morphometric data of IHC labeling of BrdU, HuCD, and GS was carried out using the Statistica 12 and Microsoft Excel 2010 software packages. All data were expressed as mean value ± standard deviation (mean ± SD) and analyzed with the SPSS software (version 16.0; SPSS Inc., Chicago, Illinois, United States). All variables measured in groups were compared using the Student’s *t*-test or one-way analysis of variance (ANOVA), followed by Newman-Keuls post-hoc analysis. Values at 0.01 < *P* < 0.05 and *P* < 0.01 were considered statistically significant.

## 3. Results

### 3.1. Morphological structure of the cerebellum in juvenile O. masou

The sections of the cerebellum from juvenile *O. masou* were stained with toluidine blue according to Nissl. The molecular, granular, and ganglionic layers of the cells were visualized (**Figure 1A**).

**Figure 1.**
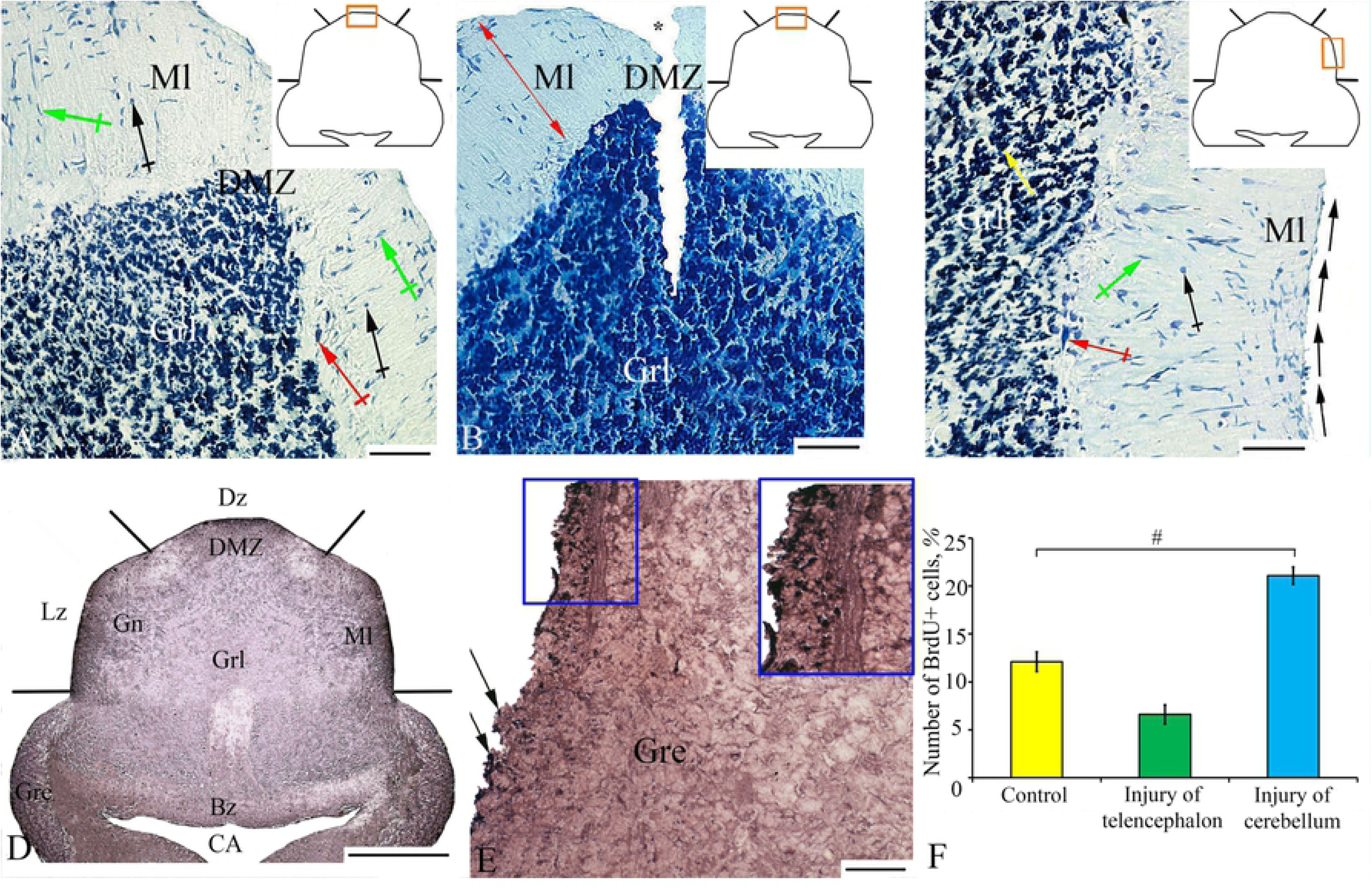
Morphological structure (A–C) and BrdU-labeling (D–F) of *Oncorhynchus masou* cerebellum. (A) Dorsal zone of intact cerebellum: figured black arrows indicate type I cells; green arrows, type II cells; red arrows, type IV cells. (B) Dorsal zone of the cerebellum at 3 days post-injury: red arrows indicate direction of radial cells migration. (C) Lateral zone of the cerebellum after injury: black arrows indicate direction of tangential migration: black arrows indicate type I cells; green arrows, type II cells; red arrows, type IV cells; and yellow arrows, type V cells. (D) General view of the cerebellum after BrdU-labeling with radial segments showing the boundaries of cerebellar zones: Dz, dorsal; DMZ, dorsal matrix; Lz, lateral; Bz, basal zone; CA, *cerebellum aqueduct*; Gre, granular eminencia. (E) Granular eminencia of cerebellum after injury: black arrows indicate of BrdU+ cells, blue rectangle outlines the BrdU+ cells aggregation. (F) Proliferative index of BrdU+ cells in the cerebellum of *Oncorhynchus masou* in the control (yellow columns), after telencephalon injury (green columns), and cerebellum injury (blue columns) (*n* = 5 in each group; error bars are SD). (A–C) Toluidine blue staining by Nissl; (D, E) BrdU-immunolabeling on transversal cerebellar sections (Dolbeare, 1995). Radial segments in the insets designate the boundaries of zones: Dz, dorsal zone; Lz, lateral; Bz, bazal zone; Ml, molecular layer; Grl, granular layer; Gn, ganglionic layer. Black asterisk indicates the area of injury; white asterisk, Purkinje cells of the ganglionic layer. Scale bars: (A–C, E) 100 µm and (D) 500 µm.

According to the studies of the cerebellum of adult trout *Salmo gairdnery*, conducted by Pouwels (1978b), we distinguished five classical neuronal cell types: the granule cells and the Golgi cells in the granular layer, the Purkinje cells and the eurydendroid cells in the ganglionic layer, and the stellate cells in the molecular layer.

To characterize cerebellum, the dorsal, lateral, and basal regions of *corpus cerebellum* (CCb) were examined (**Figure 1A**). The cell composition of all the cerebellar regions included five morphological types of cells, the morphometric parameters of which are given in **Table 1**.

**Table 1.**
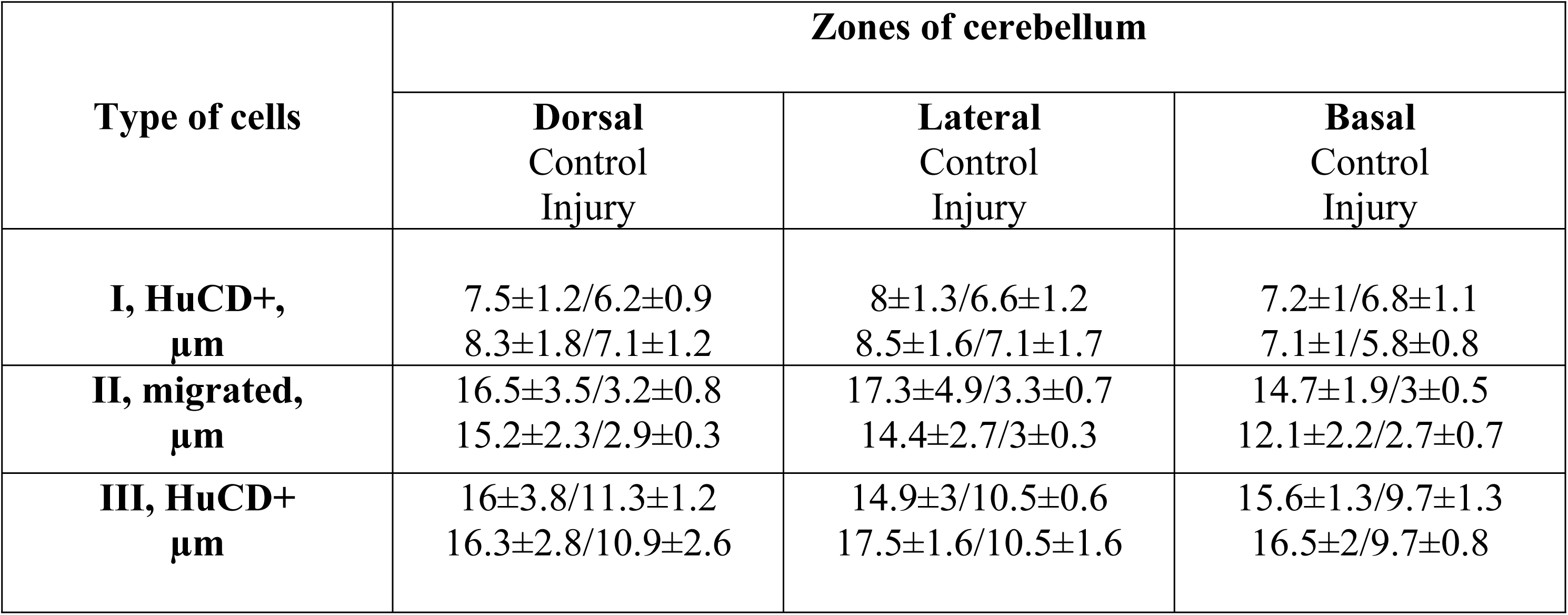
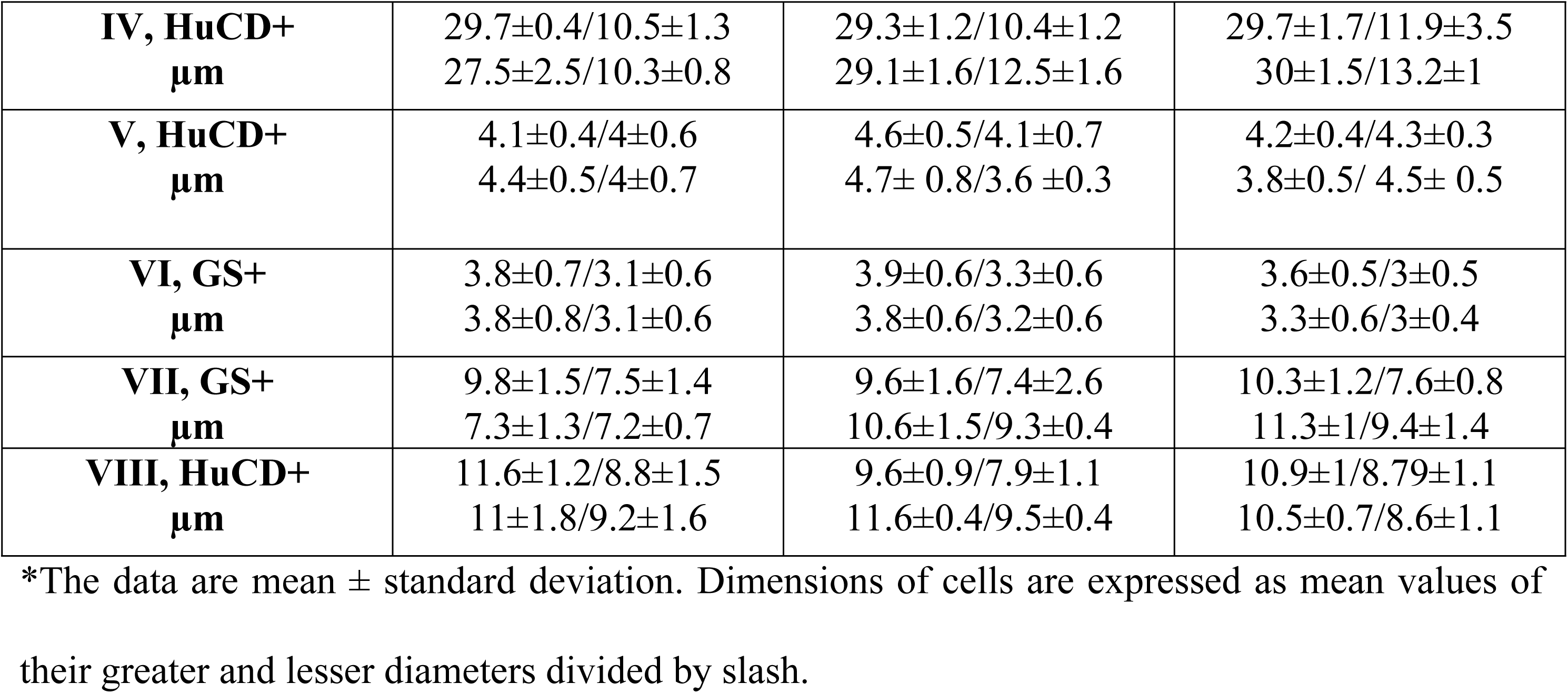
Morphometric characteristics of cells (stained by Nissl) in the dorsal, basal, and lateral zones of the cerebellum of juvenile masu salmon, *Oncorhynchus masou*, in the control *vs*. after injury (M ±SD)*.

In the molecular layer, we identified cells of a rounded/oval shape with the soma of large and small sizes. We assume that this is a single type of cells (stellate cells) at different stages of their growth, referred to as type I cells (stellate cells). In addition, we found cells which had a rod-like shape, or type II cells (**Table 1**). In the control animals, the most of the type I and type II cells were located in the middle part of the molecular layer (**Figure 1A**), where the density of distribution of such cells was rather high (**Figure 2A, D and G**). After injury, the number of type II (rod-shaped) cells in the molecular layer increased; the cells formed radially oriented, longitudinal rows (Fig. 1B). The results of the quantitative analysis showed that after the injury the number of cells in the molecular layer increased in the dorsal (**Figure 2A**) and lateral (**Figure 2D**) zones, respectively. The greatest increase in the number of cells after injury was found in the dorsal (**Figure 2A**) and lateral zones (**Figure 2D**). In the molecular layer of the basal zone, the number of cells after injury, on the contrary, decreased (**Figure 2G**).

**Figure 2.**
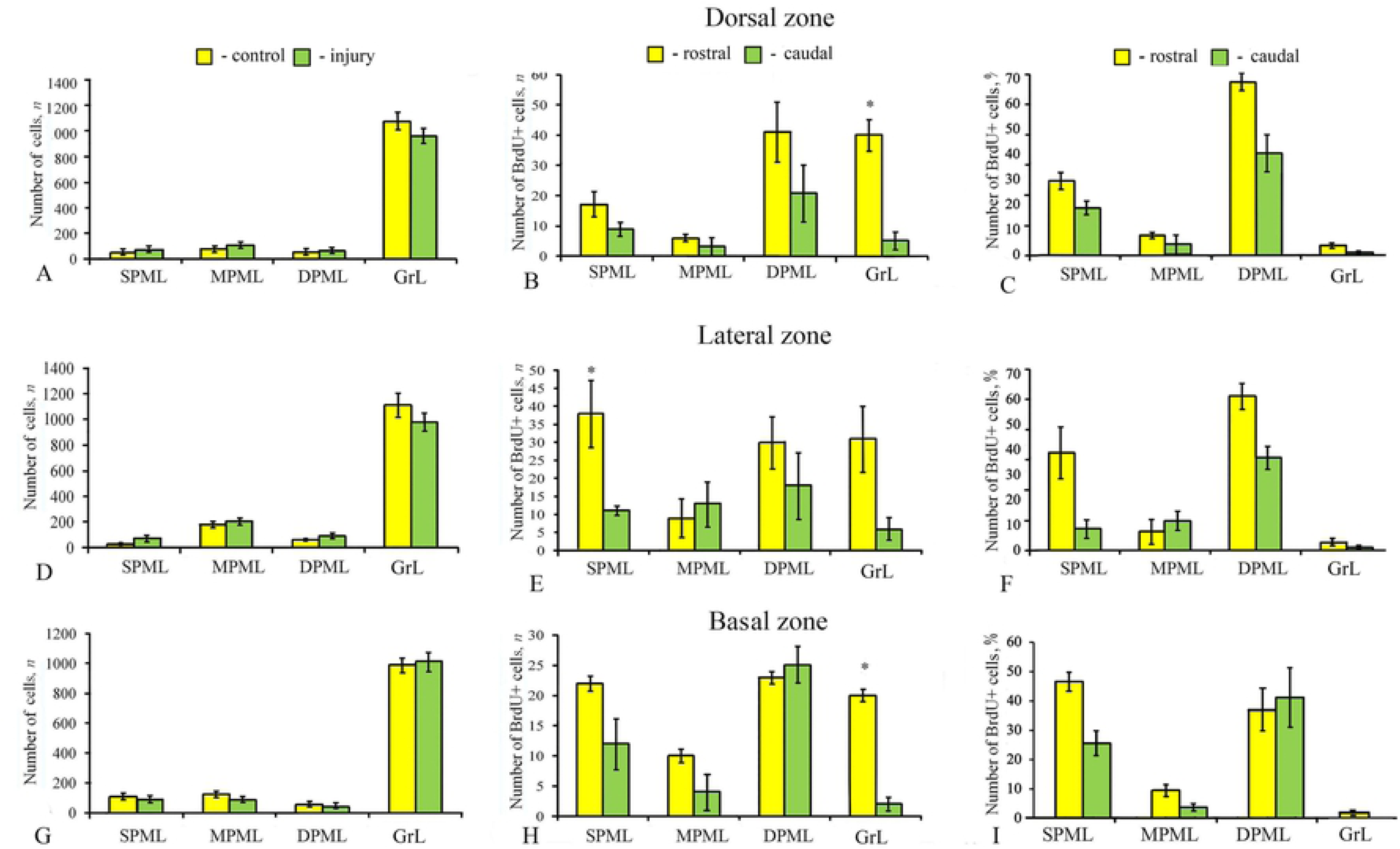
The ratio of cells stained by Nissl and BrdU-labeled cells in the cerebellum of juvenile *Oncorhynchus masou* in the control and after injury. (A, D, G) Number of cells stained by Nissl (mean ± SD): yellow columns designate areas of CCb in control animals; green columns, after injury. (B, E, H) Distribution of BrdU+ cells in the rostral and caudal parts of intact CCb: yellow columns designate the number of cells in the rostral part of CCb; green columns, the number of cells in the caudal part; error bars are SD (*n* = 5; \**P* < 0.05). (C, F, I) Proliferative index in intact CCb; SPML, surface part of the molecular layer; MPML, middle part of the molecular layer; DPML, deep part of the molecular layer; GrL, granular layer. Student’s *t*-test was used to determine significant differences between the rostral part of CCb and the caudal one.

In the ganglionic layer, the soma of pear-shaped Purkinje cells (type III) and elongated bipolar eurydendroid cells (EDC) (type IV) was stained with toluidine blue (**Table 1**).

In the surface part of the molecular layer of the lateral zone, the tangential migration of a large number of type II (rod-shaped) cells was revealed after injury (**Figure 1C**). The patterns with tangential cell migration were not detected in the control animals. In the dorsal zone near migrating type II (rod-shaped) cells, we observed the presence of type I (rounded) cells. An aggregation of several cells was considered as a cluster. Rounded cells of type I formed clusters of 3–4 elements (**Figure 1B**). In the basal and lateral zones of the cerebellum, no similar pattern of cellular distribution was detected after the injury.

In the granular layer of cerebellum, most of type V granular cells were stained with toluidine blue; the size of their soma was ca.5 μm (**Table 1**). In the dorsal and lateral zones of the granular layer, the number of cells decreased after the injury (**Figure 2A and D**). In the basal zone, the number of cells somewhat increased (**Figure 2G**). In the granular layer of the cerebellum, intensely labeled cells appeared occasionally; the size of their soma exceeded 5 μm. These single cells probably correspond to the Golgi cells of other vertebrates. In the area of white matter, single type V cells were found.

### 3.2. Experimental BrdU labeling

After the experimental treatment of BrdU in the cerebellum of juvenile *O. masou*, we estimated the number of BrdU-labeled cells and cells’ nuclei at 3 days post-injury. As a result of BrdU-labeling, positive elements were stained dark violet, but the white matter of cerebellum, the periventricular region, and the extracellular substance of the molecular layer were immune negative zones (**Figure 1D**). In the control animals, BrdU+ cells and, in some cases, BrdU+ nuclei were identified in the molecular and granular layers of the dorsal, lateral, and basal regions. According to the classification of Trainello (Trainello et al., 2014), we differentiated two types of BrdU-labeled elements: cells’ nuclei with a diameter of ca. 3.5 μm and cells with a soma diameter of 5 μm. Cells and nuclei labeled with BrdU were detected in the superficial part of molecular layer (SPML), in the middle part of the molecular layer (MPML), and in the deep part of the molecular layer (DPML), and also in the granular layer (GrL).

In the DPML of the dorsal, basal, and lateral zones of cerebellum, small groups of negative Purkinje cells, surrounded by 2–3 BrdU+ cells (nuclei) of 3.5–5 μm in size, were detected in the control animals and in both types of injury (cerebellar and telencephalic, respectively). The largest aggregation of BrdU+ cells after injury was detected in the DPML of the dorsal, lateral, and basal zones and in *granular eminentia* (**Figure 1E**).

#### 3.2.1. Distribution of BrdU+ cells in the rostral part of intact cerebellum in juvenile O. masou

The morphometric analysis of data showed similar cell sizes from different zones of the rostral and caudal parts of cerebellum both in the control animals and after the mechanical injury (**Tables 2 and 3**). In the rostral part of cerebellum, in the dorsal zone, the greatest number of BrdU+ cells was detected in the DPML and in the GrL; the smallest number, in MPML (**Figure 2B**). In the lateral zone, the maximum number of cells was detected in the SPML (*P* < 0.05, **Figure 2E**); a smaller number, in the DPML. The number of cells in the MPML was minimum (Figs. 2E; 3B). In the basal zone, the number of BrdU+ cells in the SPMLs, in the DPML, and in the GrLs averaged at 20 ± 2 cells (*P* < 0.05, **Figure 2H**). In the MPML, the number of labeled cells decreased twofold. (**Figure 2H**). In the MPML of the dorsal and basal zones, the intensely BrdU-labeled cells had a low density of distribution. In aggregations of BrdU+ cells, the intensity of labeling was sometimes lower than that of individual cells.

**Table 2.** Morphometric characteristics of BrdU+ cells in the dorsal, basal, and lateral zones of the cerebellum of juvenile *Oncorhynchus masou* in the control *vs*. after injury (M ± SD)*

| Area of cerebellum | Dorsal zone,<br>$\mu\text{m}$ | Lateral zone,<br>$\mu\text{m}$ | Basal zone,<br>$\mu\text{m}$ |
| --- | --- | --- | --- |
| Surface part of the molecular layer<br>Control<br>Injury | 4.9±0.3/4.1±0.5<br>5.5±0.6/3.9±0.8 | 5±0.3/3.5±0.4<br>4.9±0.4/3.5±0.5 | 4.6±0.4/3.7±0.6<br>5.1±0.5/3.7±0.8 |
| Middle part of the molecular layer<br>Control<br>Injury | 4.3±0.6/3.7±0.8<br>5.5±0.6/3.9±0.8 | 4.5±0.6/3.7±0.6<br>4.9±0.4/3.5±0.5 | 4.1±0.7/4±0.8<br>5.1±0.5/3.7±0.8 |
| Deep part of the molecular layer<br>Control<br>Injury | 5±0.4/3.5±0.8<br>5.7±1.2/4.5±1 | 5.1±0.8/4.4±0.8<br>4.5±0.7/3.7±0.6 | 4.6±0.9/3.6±0.7<br>5.3±0.9/4±0.6 |
| Granular layer<br>Control<br>Injury | 4.4±0.8/3.6±0.6<br>5.4±0.9/4.5±0.7 | 4.8±0.8/3.8±0.6<br>4.6±0.9/3.5±0.5 | 3.9±0.8/3.4±0.3<br>5.2±0.7/4±0.7 |

\* The data are mean ± standard deviation. Dimensions of BrdU+ cells are expressed as mean values of their greater and lesser diameters divided by slash.

**Table 3.** Morphometric characteristics of BrdU+ cells in the dorsal, basal, and lateral zones in the caudal part of the cerebellum of juvenile *Oncorhynchus masou* in the control *vs*. after injury (M ± SD)*

| Area of cerebellum | Dorsal zone,<br>$\mu\text{m}$ | Lateral zone,<br>$\mu\text{m}$ | Basal zone,<br>$\mu\text{m}$ |
| --- | --- | --- | --- |
| Surface part of the molecular layer<br>Control<br>Injury | 5±0.2/3.5±0.8<br>4.8±0.4/4.1±0.8 | 4.8±0.3/4.1±0.5<br>5.2±0.4/3.9±0.7 | 5.2±0.3/3.7±0.7<br>5±0.4/3.5±0.8 |
| Middle part of the molecular layer<br>Control<br>Injury | 5±0.4/3.9±0.5<br>5.1±1.1/4.2±0.8 | 4.3±0.8/4±0.7<br>4.8±0.9/4±0.7 | 5.1±1/4±0.6<br>4.7±0.9/3.9±0.6 |
| Deep part of the molecular layer<br>Control<br>Injury | 4.2±0.8/3.3±0.2<br>5.3±0.8/4±0.6 | 4.6±1/4±1<br>5.1±0.8/4.3±0.7 | 4.4±0.7/3.6±0.5<br>4.6±0.9/3.8±0.7 |
| Granular layer<br>Control<br>Injury | 4±0.4/3.5±0.3<br>5±1/4.3±0.9 | 4.2±0.8/4.1±0.9<br>4.6±0.6/4±0.6 | 3.9±0.6/3.7±0.5<br>4.8±0.5/4±0.5 |

\*- The data are mean ± standard deviation. Dimensions of BrdU+ cells are expressed as mean values of their greater and lesser diameters divided by slash.

In the MPML of the lateral zones, several clusters of 3–4 cells were identified; in the basal part, a cluster of four cells. In the SPML of the lateral zone, the BrdU+ cells were distributed evenly, at a distance of 8–10 μm from each other (**Figure 3B**). In DMZ, the density of distribution of BrdU+ cells was significantly higher as compared with other areas of the molecular layer (**Figure 3B**). In the DPML and GrL of the dorsal area, clusters of 8–12 labeled cells were found (**Figure 3A**). In the basal area, such clusters were not detected. Large clusters of BrdU+ cells were observed in the GrL (**Figure 3D**).

**Figure 3.**
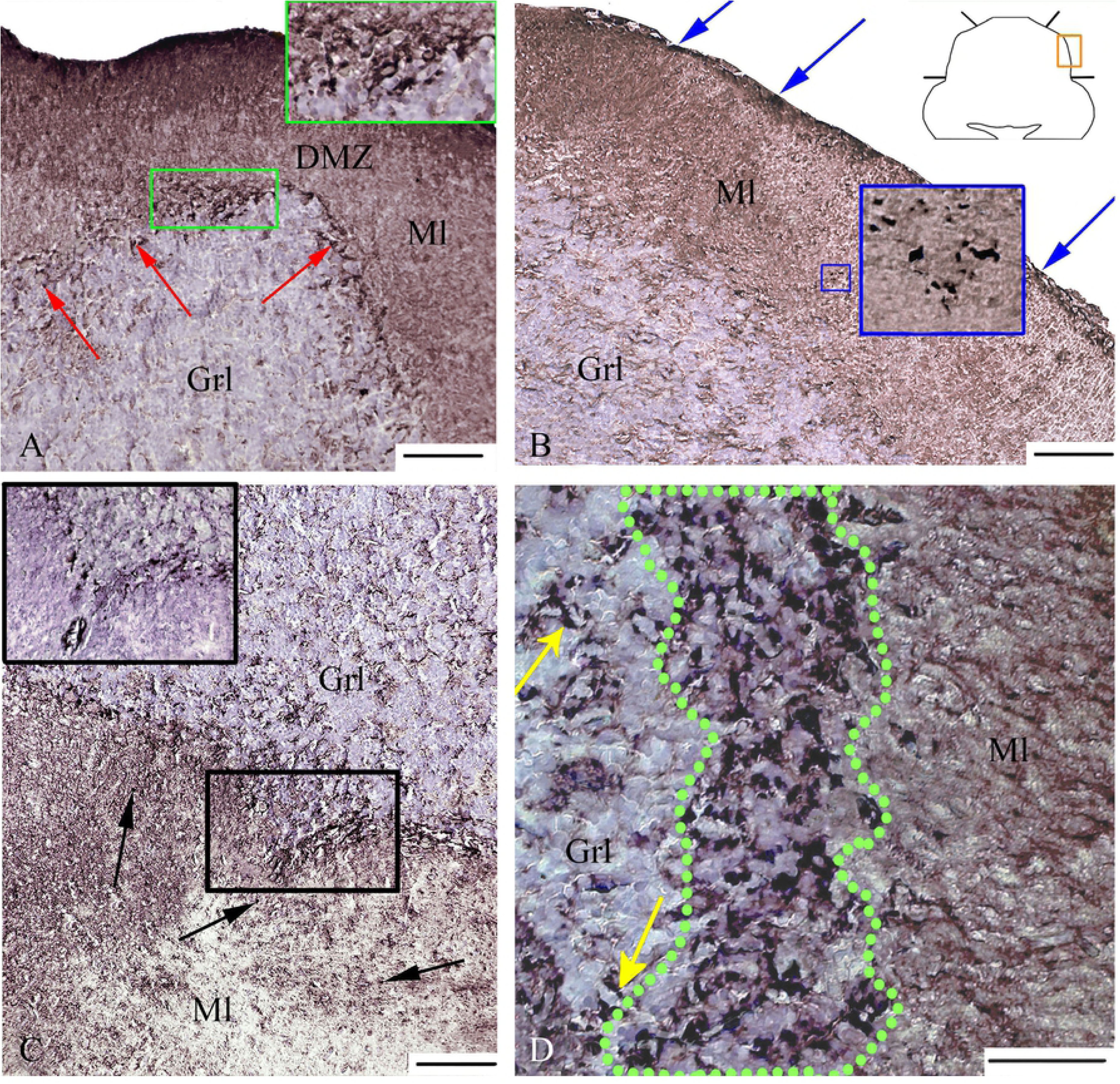
BrdU-labeling in intact cerebellum of *Oncorhynchus masou*. (A) Dorsal zone of the caudal part of cerebellum: green rectangle outlines a BrdU+ cells aggregation in the deep part of the molecular layer. (B) Lateral zone of the rostral part of cerebellum (outlined by rectangle in the inset). Blue rectangle outlines the BrdU+ cells aggregation in the middle part of the molecular layer. (C) Basal zone of the caudal part of cerebellum: black rectangle outlines the ventral tip of the basal zone. (D) Granular layer of the rostral part of cerebellum: green dotted line outlines an aggregation of BrdU+ cells. Dz, dorsal zone; Lz, lateral; Bz, basal zone. Ml, molecular layer; GrL, granular layer. Black arrows indicate BrdU+ cells in the middle part of the molecular layer; yellow arrows, BrdU+ cells in the granular layer; red arrows, an aggregation of cells in the deep part of molecular layer; blue arrows, cells at the surface of the molecular layer. BrdU-immunolabeling on transversal cerebellar sections (Dolbeare, 1995). Scale bars: (A, B, C) 100 µm and (D) 50 µm.

The DPML of the dorsal and lateral areas showed the largest proliferative index (54.2% and 58.2%); the smallest proliferative index (37.1%) was recorded from the basal zone (**Figure 2C, F and I**).

#### 3.2.2. Distribution of BrdU+ cells in the caudal part of intact cerebellum in juvenile O. masou

In the caudal part of cerebellum, the pattern of distribution of BrdU+ cells was significantly different from the rostral one. The number of labeled cells in the caudal part of the cerebellum was lower than in the rostral part; the distribution of BrdU+ cells in different layers was specific (**Figure 2B, E and H**). In the DPML of the dorsal zone, the number of labeled cells was 2 times as low as that in the rostral part of cerebellum (**Figure 2B**). In the GrL of the caudal part, the number of BrdU+ cells was significantly lower than in the rostral one (*P* < 0.05, **Figure 2B and H**). In different areas of the molecular layer, no significant differences in the number of labeled cells were found between the rostral and caudal parts of cerebellum.

In the lateral zone of the caudal part, there were separate labeled cells and small groups of cells in the MPML. The number of BrdU+ cells in the SPML was 1.7 times as low as that in the rostral part (**Figure 2E**). In the basal zone, no significant differences in the number of labeled cells in the molecular layer between the rostral and caudal parts were observed (**Figure 2H**). However, in the ventral apex of the basal zone, the number of BrdU+ cells increased distinctly and amounted to 45 ± 7 in the DPML (**Figure 3C**). In the rostral part of the corresponding zone, there were no apparent clusters of cells (**Figure 2H**).

The results of the quantitative analysis showed the maximal proliferative index in the caudal part of cerebellum: in the DPML of the dorsal and lateral zones (43% and 41% correspondently). The minimum proliferative index recorded from the GrL of the caudal part (**Figure 2C, F and I**). In general, the proliferative index of BrdU-labeled cells in the GrL of the rostral part of cerebellum was 7.5– 8.5 times as high (*P* < 0.05) as that of the caudal part (Fig. 2C, F, I). Such significant differences indicate a gradient distribution of cells in different parts of cerebellum.

#### 3.2.3 Distribution of BrdU+ cells in the cerebellum of juvenile O. masou after injury

The zone of injury extended through the dorsal surface of cerebellum, from the dorsal part of the molecular layer into the deep part of the GrL (**Figure 4A**). At 3 days post-injury, the number of BrdU+ cells around the injury zone increased, both in the rostral and caudal parts of cerebellum. In the dorsal zone of the rostral part, containing the puncture area, the number of BrdU+ cells increased significantly, especially in the SPML (**Figure 4B**). However, in the DPML, the number of labeled cells was, on the contrary, smaller than in the control (**Figure 5A**). In the dorsal zone of the caudal part, the number of cells in the molecular and granular layers increased more than in the rostral part. The number of labeled cells in the SPML and GrL was significantly greater (*P* < 0.05, **Figure 5B**).

**Figure 4.**
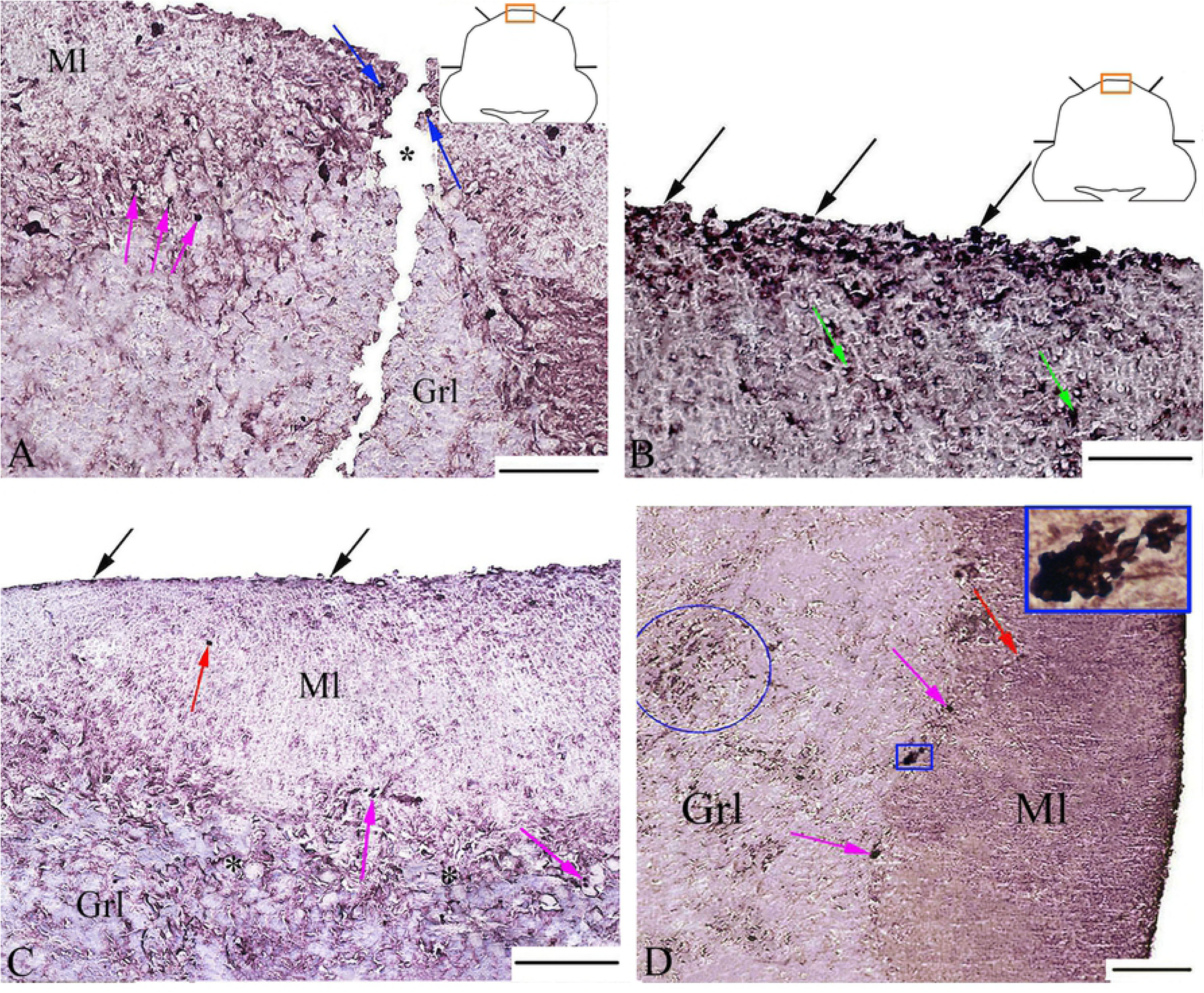
Cerebellum of *Oncorhynchus masou* after injury. (A) The area of injury in the dorsal zone of cerebellum (outlined by rectangle in the inset). (B) Dorsal zone of the rostral part of cerebellum. (C) Lateral zone of the rostral part of cerebellum. (D) Lateral zone of the caudal part of cerebellum after injury: blue oval outlines a BrdU+ cells aggregation in the granular layer; blue rectangle, a BrdU+ cells aggregation in the deep part of the molecular layer. BrdU-immunolabeling on transversal cerebellar sections (Dolbeare, 1995). Ml, molecular layer; GrL granular layer; Gn, ganglionic layer. Asterisk indicates the area of injury; black arrows, BrdU+ cells at the surface of the molecular layer; red arrows, BrdU+ cells in the middle part of the molecular layer; green arrows, single BrdU+ cells in the middle part of the molecular layer; blue arrows, cells near the area of injury; pink arrow, a cellular aggregation in the deep part of the molecular layer. Scale bars: (A, C, D) 100 µm and (B) 50 µm.

**Figure 5.**
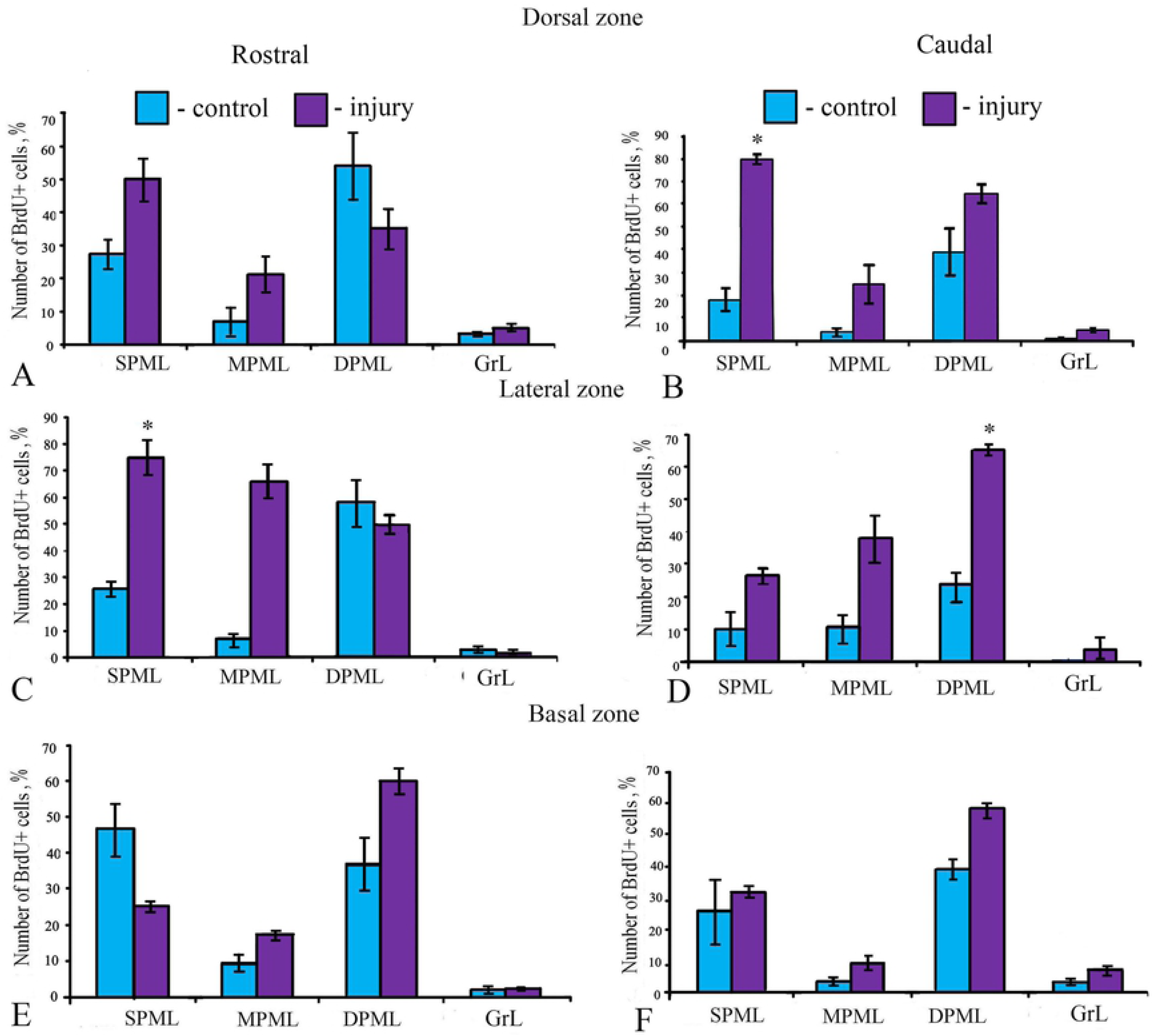
Proliferative index of BrdU+ cells in the rostral (A, C, E) and caudal (B, D, F) parts of the cerebellum of *Oncorhynchus masou* in the control and after injury. Blue columns show the number of BrdU+ cells in the control animals; purple columns, the number of BrdU+ cells after injury of cerebellum. SPML, surface part of the molecular layer; MPML, middle part of the molecular layer; DPML, deep part of the molecular layer; GrL, granular layer. Student’s *t*-test was used to determine significant differences between the control animals and those after injury (*n* = 5 in each group; \**P* < 0.05 *vs*. control group).

In the rostral part, the number of labeled cells in the molecular layer in the lateral zone increased after injury, especially in the SPML (**Figure 4C**), and in the caudal one it increased (**Figure 4D**). The most significant increase in the number of BrdU+ cells was detected in the GrL (**Figure 4C**). At the outer boundary of the molecular layer of the dorsal zone, the greatest density of distribution of BrdU+ cells was observed (**Figure 4A**); the number of cells in the lateral zone was significantly lower (**Figure 4C**).

In the basal area of the rostral part, a significant increase in the number of BrdU+ cells (P < 0.05) was detected in the middle and deep parts of the molecular layer (**Figure 5E**). In the caudal part, an increase in the number of labeled cells was found in all parts of the molecular layer after injury (**Figure 5F**). In the GrL, the number of labeled cells significantly exceeded the corresponding values in the control animals.

In the MPML of the dorsal part, located near the puncture zone, a large number of intensively BrdU+ labeled cells were found (**Figure 4A**). The greatest number of BrdU+ cells was detected in the GrL of the dorsal and lateral zones. At the boundary with the molecular layer, such cells formed small clusters (**Figure 4D**).

A quantitative analysis of the proliferative activity after injury showed an increase in the proliferative index in most parts of the rostral part, especially in the surface part, of the molecular layer (*P* < 0.05, **Figure 5A and C**) and caudal regions (**Figure 5B and D**). The most significant increase in the proliferative index was recorded from the SPML in the dorsal zone and in the DPML in lateral zone in the caudal part (*P* < 0.05, **Figure 5B and D**), which clearly indicates the predominant localization of reactive neurogenic niches in these zones. In the surface layer of the basal zone of the caudal part of the intact cerebellum, the proliferative index is higher than in the respective area after injury (**Figure 6E**). However, in other areas of the damaged cerebellum, the proliferative index increases. It is possible that the decrease in proliferative activity in the surface layer of the basal zone after injury is associated with the migration of BrdU+ cells into the deeper layers of the cerebellum to accelerate the recovery process. As a result of the study of the rostral and caudal parts of cerebellum, different patterns of proliferative activity were observed. In the intact animals, the number of BrdU+ cells in the rostral part was much larger than in the caudal one, which indicates a greater involvement of the rostral regions in constitutive neurogenesis. After injury, the number of labeled cells in the caudal part significantly exceeded that of the rostral part. We believe that this phenomenon is associated with a greater involvement of the caudal regions of cerebellum in reparative neurogenesis.

**Figure 6.**
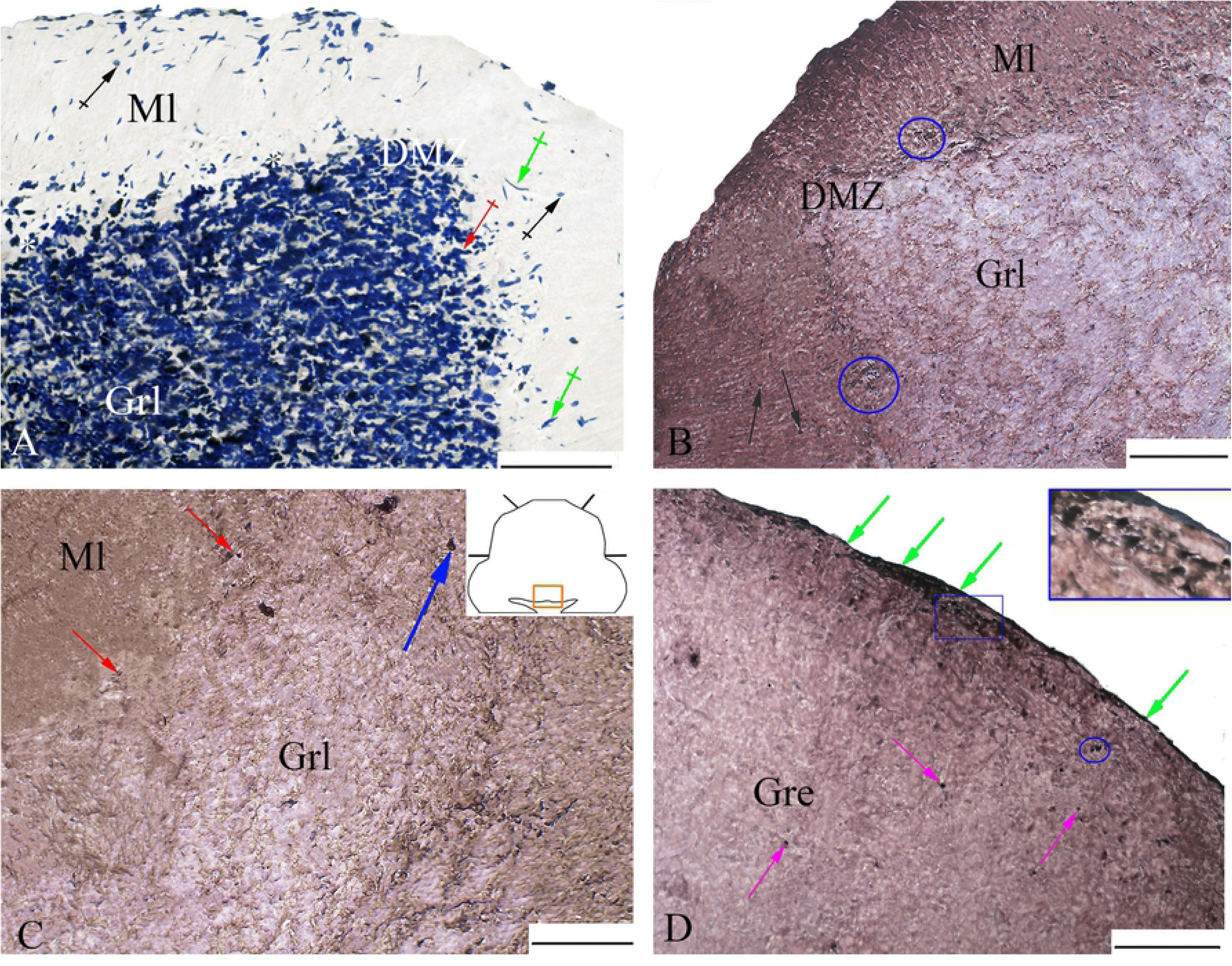
Nissl staining (A) and BrdU-labeling (B-D) in the cerebellum of juvenile *Oncorhynchus masou* at 3 days after telencephalon injury. (A) Dorsal zone. Black arrows indicate type I cells; green arrows, type II cells; red arrows, type IV cells; (B) Dorsal zone: blue ovals outline a cluster of BrdU+ cells. (C) Basal zone (outlined by rectangle in the inset). Blue arrows indicate BrdU+ cells in granular layer. (D) Granular eminencia. Green arrows indicate BrdU+ cells in the surface part of the granular eminencia; pink arrows, BrdU+ cells in the deep part of the granular eminencia; blue oval, aggregations of BrdU+ cells; blue rectangle outlines a BrdU+ cells aggregation in the surface part of the granular eminencia. (A) Toluidine blue staining by Nissl (B–D), BrdU-immunolabeling on transversal cerebellar sections (Dolbeare, 1995). Dz, dorsal zone; Lz, lateral zone; Bz, basal zone; Ml, molecular layer; GrL, granular layer. Black arrows indicate BrdU+ cells in the middle part of the molecular layer; red arrows, aggregations of cells in the deep part of the molecular layer. BrdU-immunolabeling on transversal cerebellar sections (Dolbeare, 1995). Scale bars: (A–D) 100 µm.

#### 3.2.4 Distribution of BrdU+ cells in cerebellum of juvenile O. masou after the telencephalon injury

On day 3 after the telencephalon injury, the changes in the patterns of proliferative activity in the different zones of cerebellum became obvious. A general view of the cerebellum after the telencephalon injury, stained with toluidine blue according to Nissl, is shown in Fig. 6A. In the dorsal, lateral, and basal zones, intensely BrdU-labeled cells in the DPML formed clusters of 2–3 elements (**Figure 6B**). The greatest number of labeled cells was found in the dorsal zone (**Table 4**). The cells were evenly distributed at the outer boundary of the molecular layer; single BrdU+ cells were encountered in the MPML (**Figure 6B**). In the basal zone, the highest density of moderately BrdU-labeled cells in the DPML was detected at the ventral apex. Numerous aggregations containing up to 10 cells were identified in the GrL of the dorsal zone after the telencephalon injury, as it was in the similar region after the damage to the cerebellum. In the molecular layer of the basal zone, there were no BrdU+ cells in the middle part (**Figure 6C**).

**Table 4.** Morphometric characteristics of BrdU+ cells in the dorsal, basal, and lateral zones of the cerebellum of juvenile *Oncorhynchus masou* in the control *vs*. after telencephalon injury (M ± SD)*

| Area of cerebellum | Dorsal zone,<br>$\mu\text{m}$ | Lateral zone,<br>$\mu\text{m}$ | Basal zone,<br>$\mu\text{m}$ |
| --- | --- | --- | --- |
| Surface part of the molecular layer | 4.7±0.6/3.5±0.5 | 4.9±0.5/3.9±0.6 | 5.1±0.5/3.7±0.4 |
| Middle part of the molecular layer | 5.1±1/4.2±0.7 | 4.6±1/3.9±1.3 | 5±1/3.9±0.7 |
| Deep part of the molecular layer | 5.1±0.9/3.6±0.3 | 4.3±0.8/4±0.7 | 5.1±1/4±0.6 |
| Granular layer | 5.1±0.4/4.7±0.6 | 3.9±0.6/3.7±0.5 | 4.1±0.7/3.8±0.6 |

\*- The data are mean ± standard deviation. Dimensions of BrdU+ cells are expressed as mean values of their greater and lesser diameters divided by slash.

An analysis of the quantitative distribution of BrdU+ cells in the dorsal zone after the telencephalon injury showed a decrease in the number of labeled cells in the molecular and granular layers, and the total number of proliferating cells after the damage to the brain was lower as compared to the cerebellar injury (**Figure 7A**). A similar trend was observed in the lateral zone, except for the DPML (**Figure 7B**): the number of labeled cells after the telencephalon injury decreased slightly compared to that in the intact animals. An analysis of the ratio of proliferating cells in the dorsal and lateral parts after the cerebellum and telencephalon injuries also showed that the total number of proliferating cells after the cerebellum injury is higher than that after the telencephalon one (*P* < 0.05, **Figure 7A and B**). In the basal area, an increase in the number of BrdU+ cells in the surface layer and in the DPML was revealed (**Figure 7C**). In the MPML of the basal zone, labeled cells were not detected after the telencephalon injury (**Figure 7B**). The comparative characterization of BrdU+ cells in the dorsal and lateral regions after the cerebellum and telencephalon injuries showed a similar pattern (**Figure 7A and B**).

**Figure 7.**
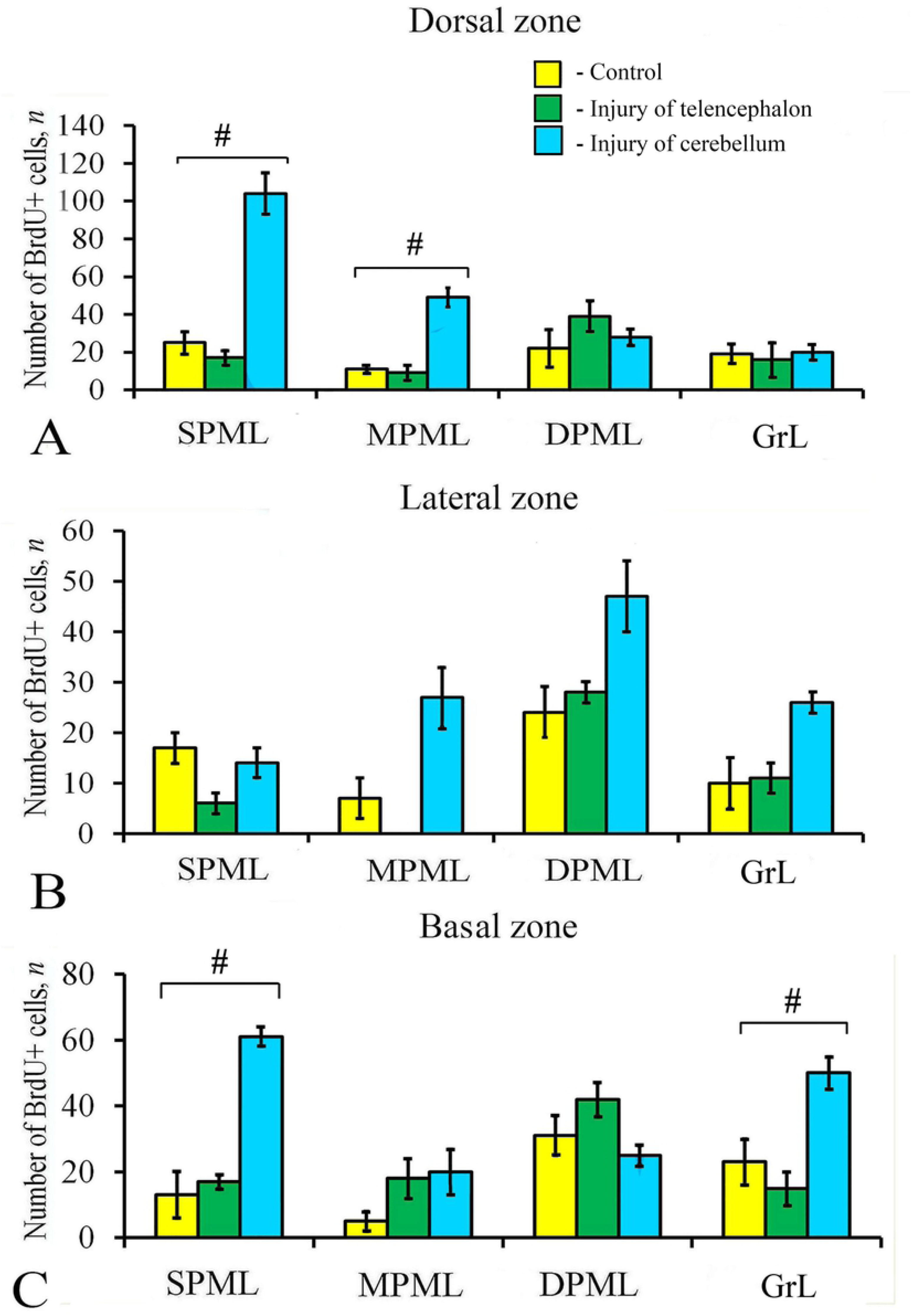
The ratio of BrdU+ cells in the cerebellum of *Oncorhynchus masou* in the control (yellow columns), at 3 days after telencephalon injury (green columns) and cerebellum injury (blue columns), (*n* = 5 in each group; error bars are SD). Yellow columns show the number of BrdU+ cells in the control animals; green columns, the number of BrdU+ cells after the telencephalon injury; blue columns, the number of BrdU+ cells after the cerebellum injury. SPML, surface part of the molecular layer; MPML, middle part of the molecular layer; DPML, deep part of the molecular layer; GrL, granular layer. Student’s *t*-test was used to determine significant differences between the control animals and those after injury (n = 5 in each group; error bars are SD).

#### 3.2.5. Distribution of BrdU+ cells in the area of granular eminences in juvenile O. masou

The largest number of BrdU+ cells in the intact and in the injured cerebellum was found in granular eminences (**Figure 1E**). The zone of granular eminences is an area of maximum cellular density and is traditionally regarded as zone of embryonic neurogenesis (Volkmann et al., 2008). In the intact cerebellum, the number of BrdU+ cells in the zone of granular eminences was on average 184 ± 28. Most of the cells were located at the surface; such cells formed intensely BrdU-labeled clusters. Similar cell clusters were also detected in the MPML; in some cases, single, moderately BrdU-labeled cells were detected. After injury, the number of BrdU+ cells was two-fold higher (*P* < 0.05), which indicates that the area of granular eminences is actively involved in reparative neurogenesis. For BrdU+ cells at the surface of granular eminences, cells in the intact and in the injured cerebellum were located at a small distance, 4–6 μm, from each other. As in the intact cerebellum, the highest activity of BrdU+ cells, forming clusters of 6–10 cells each, was characteristic of the surface of granular eminences. A significant increase in the number of cells after injury we associate, in particular, with both morphogenetic activity and the damaging effects. The results of the quantitative analysis showed that, in spite of a significant increase in the number of BrdU+ cells in the area of granular eminencies, the proliferative index was similar to those in the body of the cerebellum: 12.1 ± 1.8% in the intact cerebellum and 21.1 ± 2.7% after injury (**Figure 1F**).

After the telencephalon injury, the number of BrdU+ cells was 2.6 times as low as that in the intact cerebellum and 4.5 times as low as that in the injured cerebellum. Intensely labeled BrdU+ cells were located at a small distance from each other at the surface of granular eminences, in most cases forming intensely labeled clusters.

### 3.3. Distribution of GS+ cells in cerebellum of intact juvenile O. masou

Distribution of GS+ elements in the dorsal zone of cerebellum in juvenile *O. masou* is shown in **Figure 8A**. The GS+ cells and fibers were found in the dorsal, lateral and basal areas of CCb. In the molecular layer of the control animals, a GS activity was detected in type the VII and VII cells. Rounded type VII cells were intensely labeled; their sizes are given in Table 1. Type VI cells had the rounded soma, but their size was significantly smaller (**Table 1**). Most of them were located singly or, in some cases, grouped into two elements in the MPML or, sometimes, near the end of GS+ fibers. In the molecular layer, the distribution density of GS+ elements was higher than that in the granular layer.

**Figure 8.**
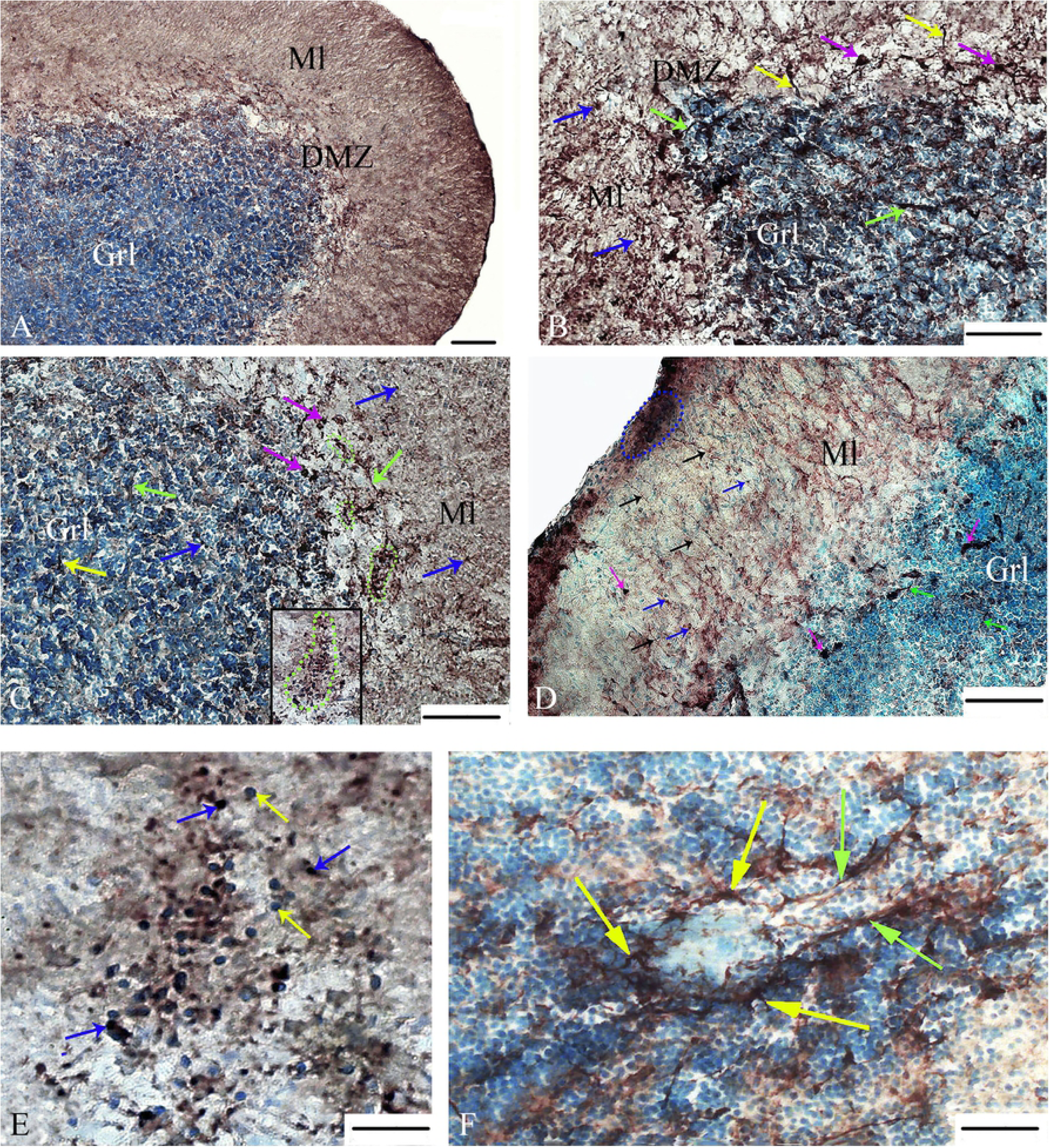
The distribution of GS in the cerebellum of intact juveniles of *Oncorhynchus masou* in the control and at 3 days post-injury. (A) General view of distribution of GS in the intact dorsal zone. (B) dorsal zone of cerebellum after injury: blue arrows, GS+ type VI cells, pink arrows, GS+ type VII cells; green arrows indicate thin, weakly labeled fibers; yellow arrows indicate thick, intensely labeled fibers. (C) Parenchymal neurogenic niches in the dorsal zone after injury: green dotted line outlines reactive parenchymal neurogenic niches containing GS+ and GS– cells (for other designations, see fig. B). (D) GS in the lateral zone of intact cerebellum: black arrows indicate GS+ fibers of radial glia; blue dotted line outlines neuroepithelial constitutive neurogenic niches containing GS+ cells. (E) Neurogenic niche in the molecular layer of the dorsal zone after injury: blue arrows indicate GS+ cells; yellow arrows, GS– cells. (F) GS+ fibers surrounding the cerebellar glomerulus. Ml, molecular layer; Grl, granular layer; DMZ, dorsal matrix zone. GS-immunolabeling on transversal cerebellar sections. Scale bars: (A–D) 100 µm and (F, E) 50 µm.

Intensively labeled, large type VII cells were detected in DMZ; the proportion of such cells per view field in the molecular layer was 51%. Type VI cells had a less intense GS-labeling; their proportion in the molecular layer was 35%.

Another type of GS+ structures in the molecular layer was represented by labeled fibers. The features of microsculpture make it possible to discriminate thinner smooth fibers (**Figure 8A**) and thick fibers with varicose thickenings as *button ent passan*. During the morphometric analysis, 5 ± 2 fragments of fibers per view field were found in the control animals (**Figure 9A**). Type VII cells in the intact animals were located in the molecular layer, infraganglionic plexus, and the granular layer. The number of their fibers accounted for 5.1% in the dorsal, 4% in the lateral, and 3.6% in the basal zones, respectively. The highest number of GS+ elements was found in the molecular layer, composed of intensely labeled type I cells, in the control animals: 45.8% in the dorsal, 37.5% in lateral, and 32.6% in the basal zones, respectively. Thin and long fibers of radial glia were found only in the lateral part of molecular layer (**Figure 8D**). Their number was 78 ± 25 elements per view field. The immunolabeling of fragments of GS+ fibers revealed 68 ± 16 per view field (**Figure 9F**). In other areas of CCb, such structures were absent. The largest part of GS+ cells was localized in the GrL, but GS+ fibers were also identified. The thin fibers were terminated with the specific end apparatus, encompassing individual granular cells or their small clusters (**Figure 8D**).

**Figure 9.**
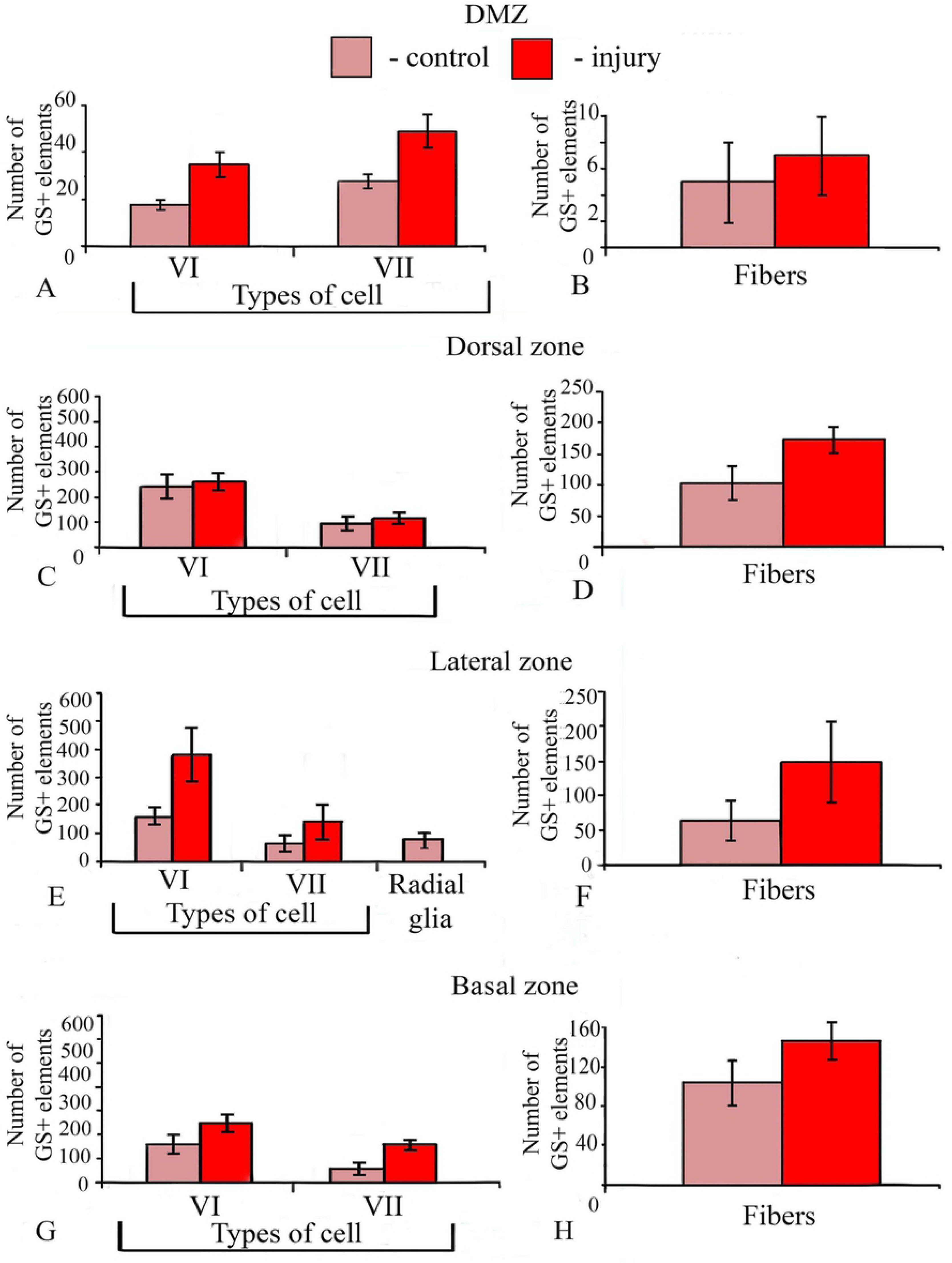
The ratio of GS+ cells and fibers in the cerebellum of intact juvenile *Oncorhynchus masou* and at 2 days post-injury: (A and B) in the dorsal matrix zone (DMZ); (C and D) in the dorsal zone; (E and F) in the lateral zone; (G and H) in the basal zone. Pink columns show the number of cells in the control animals; red columns, the number of cells at 2 days post-injury. Student’s *t*-test was used to determine significant differences between the control animals and those after injury (*n* = 5 in each group; error bars are SD).

In the GrL of the central part, we found local regions with an increased cellular density (**Figure 8F**). Such areas in CCb had a structured organization. In these regions, there was a high concentration of immunonegative cells of the GrL; the density of distribution of GS+ cells decreased towards the periphery. Areas of high cell density containing glomerular-like complexes were surrounded by more sparse zones, where type VII cells with moderate GS activity dominated (**Figure 8F**). However, the clusters of GS+ cells surrounded by fibers were characteristic mostly for zones with high cell density.

### 3.4. Distribution of GS in the cerebellum of juvenile O. masou after injury

On day 3 after the cerebellum injury in *O. masou* juveniles, we observed significant changes in the GS activity in different types of cells and fibers, both in the region adjacent to the damage zone and in more remote areas of the molecular and granular layers. The general view of cerebellum after injury is shown in **Figure 8B**. Thick GS+ fibers with varicose thickenings were oriented from granular layer to the DMZ (**Figure 8B**). After injury, the number of type VII cells and fibers (**Figure 9A–G**) gradually increased. The study of the cellular composition in the area of injury showed a 1.5-fold increase in the number of intensely labeled, large type VII cells as compared with the control. The average size of such cells on day 3 post-injury is shown in **Table 1**.

After the injury, the number of intensively labeled GS+ cells of type VII in the dorsal, lateral, and basal zones increased 1.9-fold as compared with the control (**Figure 9A, C, E and G**). Type VI cells with moderate GS activity were located singly, but more intensely labeled type VI cells formed clusters (**Figure 8B**). The appearance of new GS+ cells and their aggregations after injury was more characteristic for molecular layer (**Figure 8B**). Thus, a substantial change in the cellular composition of GS+ elements in the molecular layer and a general increase in the number of GS+ cells (1.6-fold) and GS+ fibers (1.2-fold) were observed after injury. The number of GS+ cells on day 3 after the traumatic injury gradually increased. The vast majority of GS+ cells were in the GrL. The ratio of GS+ cells in the control and injured fish was 1: 2.3 in the dorsal zone; 1 : 2 in the basal zone; and 1 : 1.9 in the lateral zone (**Figure 9A, C, E and G**). After the injury, local aggregations of small GS+ undifferentiated cells with an area of 716 ± 165 μm^2^ and large clusters with an area of 2368 ± 133 μm^2^ were detected in the dorsal part. These cell clusters were surrounded by a few heterogeneous immunopositive fibers (**Figure 8C**).

### 3.5. Infraganglionic plexus (IFGP)

IFGP is a transitional zone between the molecular and granular layer, containing bodies of Purkinje cells (PC) and bodies of eurydendroid neurons (EDC) (**Figure 8C**). This area of CCb contained both cells of the GrL and cells that occurred in the molecular layer, but most cells were represented by large ganglionic cells: PC and EDC. At IFGP of juvenile *O. masou*, the GS activity was detected in type VI and VII cells. Type VII cells in IFGP were larger in size than in cells in the molecular layer; the mean size of type VII cells was 10.3 ± 1.4 / 7.8 ± 1.1 μm in the control fish and 10.9 ± 2.1 / 8.3 ± 1.6 μm on day 3 post-injury. We found also singly distributed moderately labeled type VI cells, sometimes creating clusters (**Figure 8C**). Type VI cells in the IFGP were usually intensely immunolabeled with GS. Such cells formed small clusters consisting of 3–5 elements (**Figure 9C**). There were clusters containing both type VI and type VII cells. In all cases, the intensity of GS-labeling in such clusters was high. Clusters of small GS+ cells usually were associated with large GS+ ganglion cells (**Figure 8C**). We also revealed a neurogenic niche in the molecular layer of the dorsal zone after injury, containing a large number of GS+ cells (**Figure 8E**).

### 3.6. Enzyme immunoassay (ELISA) of GS content in O. masou cerebellum after injury

The results of the enzyme immunoassay analysis of quantitative level of GS in *O. masou* cerebellum indicate variations in the GS concentration between different time periods post-injury. Thus, in the control animals, the GS concentration was 15.7 ± 2.6 pg/mL; at 1 hour post-injury, on average 21.8 ± 1.8 pg/mL (**Figure 10**). During the first 24 hours post-injury, the maximum GS concentration was recorded at 3 h, 23.4 ± 1.9 pg/mL, and reached the control level at 12 h, 15.8 ± 3.6 pg/mL (**Figure 10**). Further observations showed that at 1 day after the cerebellum injury, the concentration of GS was 18.5 ± 2.3 pg/mL; at 2 day, it increased to 22.9 ± 0.55 pg/mL (**Figure 10**). On day 3 post-injury, the GS level again decreased to 16 ± 2.2 pg/mL and reached a maximum level of 24.2 ± 0.6 pg/mL by day 5 (**Figure 10**). Thus, the ELISA data indicate a multiple change: a decrease and increase in the GS concentration after the traumatic damage to the cerebellum of *O. masou*.

**Figure 10.**
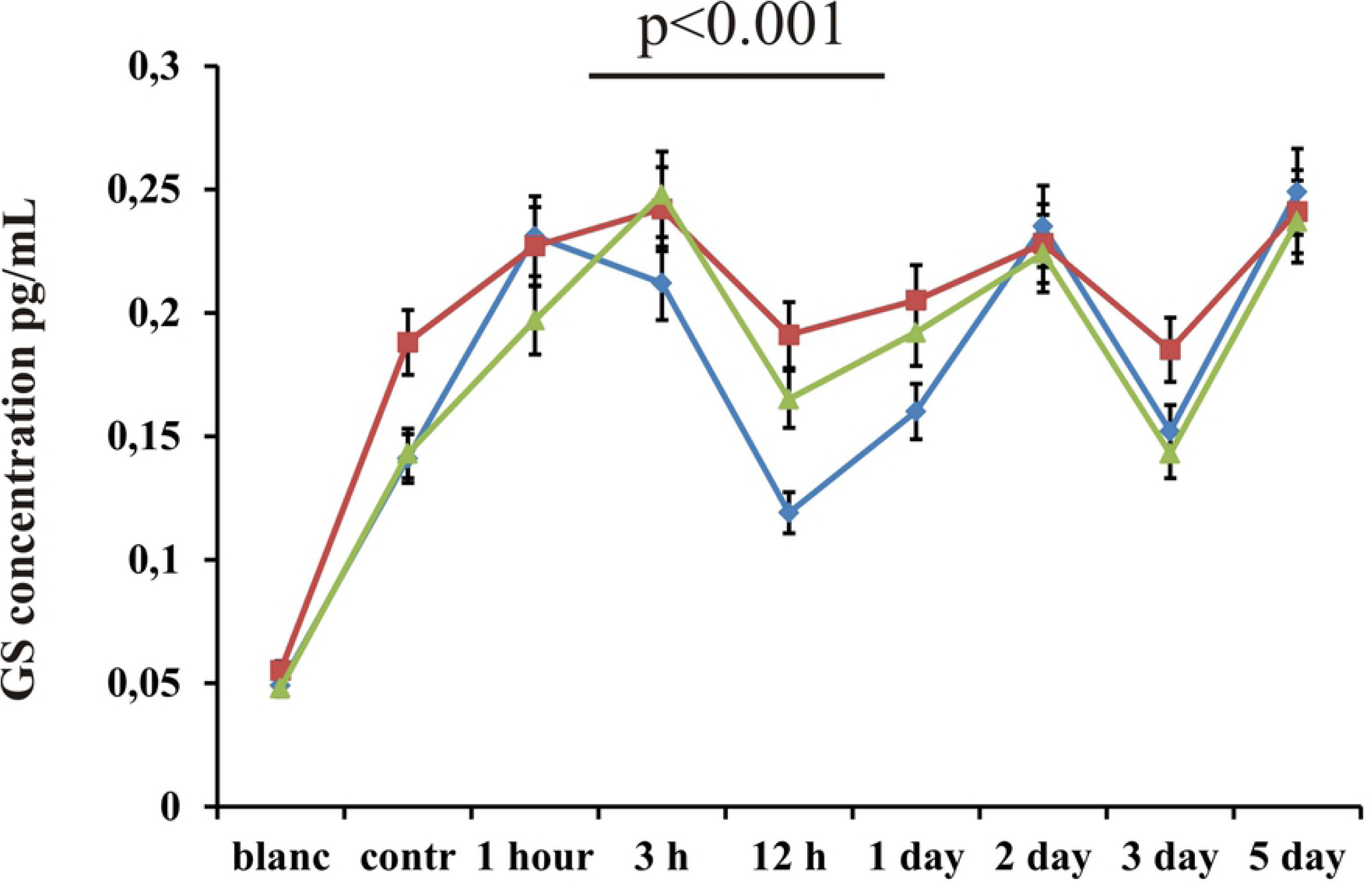
Enzyme immunoassay (ELISA) of GS concentration in the *O. masou* cerebellum after injury. The immunoassay data of glutamine synthetase concentration in the cerebellum of *Oncorhynchus masou* at different time intervals after injury. X-axis, time after injury; Y-axis, mean optical density (absorbance at 450 nm). Data are mean ± SD (*n* = 5 in each group).

### 3.7. Distribution of neuronal marker HuCD in the cerebellum of juvenile O. masou in control and after injury

HuCD+ cells were detected both at the surface of molecular layer, and in the middle part of the dorsal, lateral, and basal zones of CCb in the control fish and after the injury. For the surface layers, the presence of undifferentiated cells was constant, while the neurons in the middle part were at different stages of differentiation. We identified four types of HuCD+ cells depending on the intensity of immunolabeling, morphometric parameters, and presence/absence of processes (**Table 1**).

In the intact cerebellum, the largest number of HuCD+ cells was detected in the dorsal zone, including DMZ (**Figure 11A**). Undifferentiated, rounded type I cells dominated the area of DMZ in the control fish; such cells were weakly labeled and devoid of processes. They formed small surface clusters of 7–10 cells; in the MPML, they were located mostly separately, at a short distance from each other. The quantitative analysis showed that the proportion of such cells was 42.3% in the dorsal, 18% in the lateral, and 31% in the basal zones. In the dorsal zone, intensely labeled, rounded type VIII cells had clearly visible processes; their number was significantly larger that of type I cells. In the lateral zone, the overall density of cell distribution was similar to that in the dorsal one. However, in the lateral zone, rounded type VIII cells dominated (**Figure 11B**). The rounded and oval, intensely labeled HuCD+ cells in rare cases formed clusters of two cells, often located singly, at a short distance from each other (**Figure 12C**). In the dorsal zone, the largest number of HuCD+ cells was located near DMZ. The number of cells decreased in the lateral direction; in the dorso-lateral zone, their number was minimum (**Figure 12A**).

**Figure 11.**
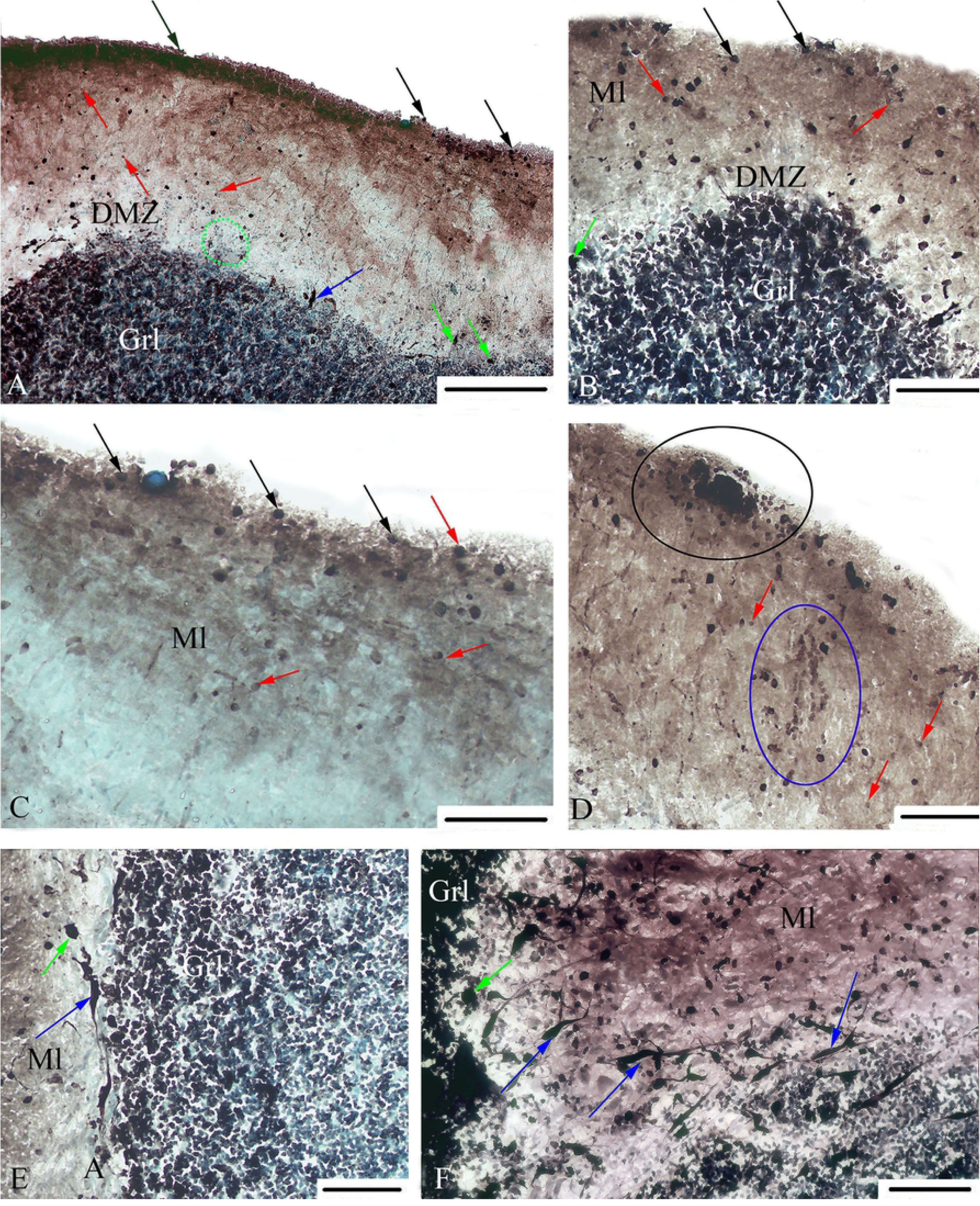
The distribution of HuCD in the cerebellum of intact juvenile *Oncorhynchus masou* and at 3 days post-injury. (A) The dorsal zone of intact cerebellum: green ovals outline parenchymal neurogenic niches containing HuCD– cells. (B) The dorsal zone of damaged cerebellum. (C) Molecular layer of the dorsal zone of intact cerebellum. (D) Molecular layer of the dorsal zone at 3 days post-injury: black oval outlines a surface neurogenic niche containing HuCD+ cells; blue oval, HuCD– migrating cells. (E) Eurydendroid (EDC) and Purkinje (PC) cells in the lateral zone at 3 days post-injury. (F) EDC in the basal zone at 2 days after injury. DMZ, dorsal matrix zone; Ml, molecular layer; GrL, granular layer. Red arrows indicate undifferentiated cells; black arrows, type I cells; pink arrows, cellular aggregations; blue arrows, EDC; green arrows, Purkinje cells. HuCD-immunolabeling on transversal cerebellar sections. Scale bars: (B–F) 50 µm and (A) 100 µm.

**Figure 12.**
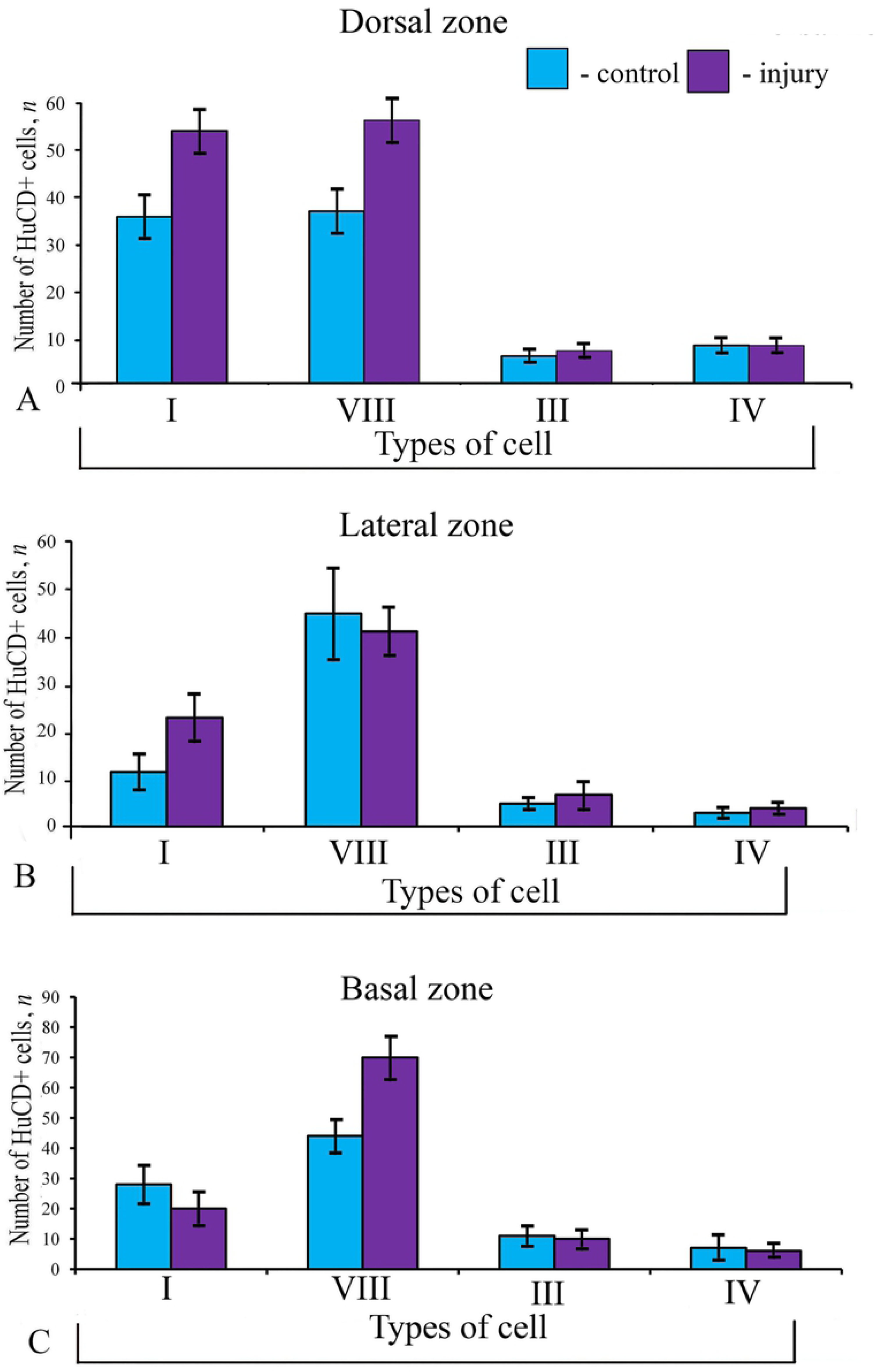
The ratio of HuCD+ cells in the cerebellum of intact juvenile *Oncorhynchus masou* and at 2 days post-injury: (A) in the dorsal zone; (B) in the lateral zone; (C) in the basal zone. Blue columns show the number of cells in the control animals; purple columns, the number of cells at 2 days post-injury. Student’s *t*-test was used to determine significant differences between the control animals and those after injury (*n* = 5 in each group; error bars are SD).

In the intact cerebellum, the pear-shaped Purkinje cells were detected in the ganglionic layer (**Table 1**). The greatest number of such cells was found in the basal zone (**Figure 11C**). Pear-shaped cells were labeled in the proximal areas of dendrites. Intensely labeled EDC were found in the dorsal, lateral, and basal zones (**Table 1**). Their number was significantly lower than in other cell forms: 6.5% in the dorsal, 7.5% in the lateral, 12.3% and in the basal zones.

In the area of the lateral and basal zones of intact cerebellum in surface layer, there were constitutive neurogenic niches containing intensely labeled, rounded type VIII cells (**Figure 11C**). At 2 days post-injury, the density of cells distribution in the DPML increased significantly (**Figure 11A and C**). The greatest concentration of cells was detected in the dorsal zone of CCb, including DMZ. The cellular composition of niches in the injured cerebellum was more heterogeneous and contained cells of various types, including undifferentiated ones than that of constitutive neurogenic niches in intact cerebellum (**Figure 11D**).

Thus, the dorsal zone of CCb exerted the most pronounced influence on reparative neurogenesis, and can also be considered as a zone where intensive processes of constitutive neurogenesis occur (**Figure 12B**). In the dorsal zone of intact cerebellum, the greatest concentration of cells was detected in the DMZ; it gradually decreased with the distance from this region. In the post-injury period, a large number of HuCD+ cells were observed throughout the dorsal zone. If in the control fish, rounded type I cells forming constitutive neuroepithelial neurogenic niches were found in the surface of lateral and basal zones; after the injury, HuCD+ cells were detected in the reactive neurogenic domains containing heterogeneous cell populations of all types, including large, intensely labeled, elongated cells. Another important observation is the increase in the total number of reactive neurogenic domains in the cerebellum. The above facts indicate the reactivation of constitutive neurogenic niches after injury. Near the neurogenic niches, radially migrating, weakly labeled cells were identified (**Figure 11D**). After the injury, in the DPML, the number of undifferentiated immunonegative cells increased 1.5-fold in the dorsal and 2-fold in the lateral parts; in the basal part, on the contrary, it decreased by a third (**Figure 12A, B and C**).

In the basal area, intensely labeled undifferentiated type 1 cells were detected in some cases, but they were not found in the other zones. In the basal area, the greatest number of rounded type VIII cells was recorded after the injury (**Figure 12C and 11F**). Large type IV cells were detected in the deep layers of all zones of the cerebellum (**Figure 11E**).The number of intensely labeled, rounded type VIII cells, sometimes with processes, was significantly larger after the injury. The largest number of type IV and V cells was detected in the basal zone (**Figure 11E**).

## 4. Discussion

### 4.1 Morphological structure of fish cerebellum

The cellular organization of the *corpus cerebelli* in teleosts is known from a few Golgi (Nieuwenhuys and Nicholson, 1969; Pouwels, 1978a-c; Murakami and Morita, 1987; Folgueira et al., 2006) and electron microscopic studies (Kaiserman-Abramof and Palay, 1969; Pouwels, 1978b). Teleosts have no cerebellar nuclei, but show a group of specialized cells, the eurydendroid cells, that are intermingled with Purkinje cells and send their axons out of the cerebellum carrying cerebellar outputs (Torres et al., 1992; Ikenaga et al., 2005; Folgueira et al., 2006).

The molecular layer contains parallel fibers, Purkinje cell dendrites, efferent cell dendrites, and stellate cells. Nieuwenhuys et al. (1974) found that the ganglionic layer of cerebellum in mormyrids contains two types of neurons, Purkinje cells, and eurydendroid cells. These authors suggest that the latter elements constitute the output system, in this respect being comparable to the cells of the deep cerebellar nuclei of higher vertebrates.

The granule layer, beneath the ganglionic layer, contains the granule cells, mossy fiber terminals, and Golgi cells. The latter are distributed throughout the granule layer (Nieuwenhuys and Nicholson, 1969; Meek and Nieuwenhuys, 1991). Granular cells give rise to parallel fibers that run in the cerebellar molecular layer and the cerebellar crest overlying the rhombencephalic lateral line region (Bass, 1982; Meek, 1992).

### 4.2. Proliferative activity in the cerebellum of juvenile O. masou after injury

BrdU-labeling in the cerebellum of juvenile *O. masou* revealed two types of elements: small rounded cells (type V cells) and nuclei of cells with a diameter of 3.5 μm. According to data of Zupanc and Ott (1999), BrdU labels a limited number of cells in the S-phase of the cell cycle. The results of studies on *O. masou* cerebellum indicate that the population of BrdU-labeled cells is more homogeneous than the cells labeled with PCNA (Stukaneva et al, 2017). As the studies on trout (Pushchina et al., 2016) and *O. masou* (Stukaneva et al., 2017) has shown, the number of PCNA+ cells in the DMZ of cerebellum after a direct damage to the cerebellum (Stukaneva et al., 2017) and to the optic nerve (Pushchina et al., 2016) is much larger than the number of BrdU+ cells after a damage to the cerebellum and telencephalon of *O. masou* (present data). Also, the population of PCNA+ cells identified in DMZ is more heterogeneous than the population of cells labeled with BrdU. A labeling of proliferative nuclear antigen (PCNA) revealed additional DNA polymerase δ (Bravo and McDonald-Bravo, 1987). In damaged CCb, a greater number of cells are found in different phases of the cell cycle and represent both resting and migrating cell forms. According to data of Waseem and Lane (1990), PCNA immunolabeling has detected both nuclear and cytoplasmic localization. Thus, the patterns of proliferating cells after PCNA labeling differ significantly in their quantitative level and morphological characteristics from those after experimental BrdU labeling.

A study of the formation of new granular cells in the cerebellum of adult zebrafish, *D. rerio*, showed that NSC and precursor cells of the cerebellum during embryonic development are created in the upper lip of rhombencephalon, similarly to those in birds and rodents (Kaslin et al., 2009). Both BrdU and PCNA labeling of cells detected extensive proliferation in the molecular layer and along the midline from the cup-like protuberance of the granular layer to the molecular layer. From there, along the vimentin+/GFAP+/BLBP+ radial glial fibers, the cells migrate radially to the surface of the molecular layer, distributing laterally during further tangential migration. Moving deeper in the radial direction from the molecular layer, they differentiate into granular neurons. Kaslin and co-authors found that the migration of glial progenitors in zebrafish is similar to the migration of glial cell progenitors in the outer granular layer of rodent cerebellum (Kaslin et al., 2009). According to Sîrbulescu et al. (2014), the number of BrdU+ cells in the ventral apex of the cerebellum in *Apteronotus leptorhynchus* is much larger than in the molecular and granular layers of other cerebellar zones (Sîrbulescu et al., 2014).

The labeling of BrdU in the cerebellum of intact *O. masou* juveniles showed that, in the dorsal zone, the greatest number of BrdU+ cells was detected in the deep part of the molecular layer; the smallest number, in its middle part. Quite large clusters of BrdU-positive cells were observed in the granular layer of cerebellum, which agrees with the data obtained for zebrafish (Kaslin et al., 2009). In general, the pattern of distribution of BrdU+ cells in the cerebellum of juvenile *O. masou* encompasses DMZ, the surface layers of the molecular layer, and also, to a lesser extent, the deep and internal parts of the molecular layer. Thus, the dorsal zone of CCb exerted the most pronounced influence on the reparative neurogenesis, and can also be considered as a zone where intensive processes of constitutive neurogenesis occur. In the dorsal zone of the intact cerebellum, the greatest concentration of cells was detected in the DMZ; it gradually decreased with the distance from this region. In the post-injury period, a large number of HuCD+ cells were detected throughout the dorsal zone.

We established that BrdU+ cells in the intact animals dominate the rostral part of cerebellum and are less represented in the caudal one. By the quantitative analysis for the intact juvenile *O. masou*, the maximuml proliferative index was obtained for the DPML of DMZ (66.3%); the minimum index, for the basal area (37.1%).

After the traumatic damage to the cerebellum, on the contrary, the number of BrdU+ cells increased significantly in the caudal part; in the rostral part, the increase in the number of proliferating cells was less pronounced. A particularly significant increase in BrdU+ cells was detected in the SPML, which is associated with the appearance of reactive neurogenic niches that arise after a traumatic injury. The results of BrdU-labeling are consistent with HuCD-immunolabeling data for the *O. masou* cerebellum, since after the injury, a large number of reactive neurogenic niches were detected in the SPML.

A significant increase in the number of proliferating cells was detected in the DPML, and, in the intact animals, the number of BrdU+ cells in this region was extremely low. In the lateral zone of the cerebellum of juvenile *O. masou*, a significant increase (*p* < 0.01) in the number of BrdU+ cells was detected in the GrL, apparently corresponding to the area of *granular eminentia* of zebrafish (Grandel et al., 2006). In the basal area, a significant increase in the number of BrdU+ cells was observed in the DPML.

After the cerebellum injury in *O. masou*, we detected an uneven proliferative activity in the surface, middle, and deep parts of the molecular layer. According to Than-Trong and Bally-Cuif (2015), neuroepithelial constitutive niches in zebrafish brain have a superficial localization. In our studies, the greatest number of BrdU+ cells in the intact animals was found in the rostral part of the cerebellum that probably indicates an intensive constitutive neurogenesis in juvenile *O. masou*. After the cerebellar injury, the greatest number of BrdU+ cells was detected in the caudal part of cerebellum. We believe that the traumatic injury stimulated the reactivation of constitutive niches, which were located both in the rostral and caudal parts of cerebellum. However, the low proliferative activity of cells of the caudal part of the intact cerebellum determinates the increase in proliferative activity in this zone after trauma, but it was more evident than in the rostral part.

We found the largest number of BrdU+ cells in granular eminences. The zone of granular eminences is an area of the maximum cellular density and is traditionally regarded as the zone of embryonic neurogenesis (Volkmann et al., 2008). After the injury of the cerebellum, the number of BrdU+ cells increased significantly (two-fold, *P* < 0.05), which indicates that the area of granular eminences is actively involved in reparative neurogenesis. After the telencephalon injury, on the contrary, the number of proliferating cells was less significant, being similar to that in the intact cerebellum. Our data are in agreement with the observations of Zupanc and Ott (1999), who recorded a great number of mitoticaly active cells from the granular eminence of the cerebellum in a brown ghost fish. In contrast, in the granular eminences of zebrafish and stickleback, they were detected in small numbers (Grandel et al., 2006). Such differences in proliferative behavior in the granular eminence can probably be explained by the method of causing injury or by adaptation of one or another species of fish to environmental conditions. However, the proliferative index was similar to that in the body of the cerebellum, and we associate this with the increased cell density in the area of granular eminence.

A comparison of proliferative activity in juvenile *O. masou* showed a much greater number of BrdU+ cells after the cerebellum injury than after the telencephalon one. According to the studies on *D. rerio*, the gliosis depends on the way in which the brain has been damaged (März et al., 2011; Baumgart et al., 2012). A strong reactivity of glial cells was observed after a direct brain injury through the skull (März et al., 2011, Kishimoto et al., 2012) and was absent in zebrafish brain after a damage of brain through the nostrils (Kroehne et al., 2011; Baumgart et al., 2012).

It has been established that after a brain injury in fish the rate of cell proliferation is much higher than in the absence of damaging effects. According to Clint and Zupanc (2001), the density of GFAP+ fibers, along which new cells migrate to the site of injury, is maximal at day 8 and remains high for at least 100 days post-injury. The presence of vimentin+ fibers in *Apteronotus leptorhynchus* (Zupanc et al., 2012) and GFAP+ fibers in *O. masou* juveniles (Stukaneva et al., 2017) suggests that intermediate filaments may participate in survival and differentiation of new cells. To date, the signals that attract radial glia fibers to the trauma site and target its processes to the hemispheres of telencephalon remain unknown (Skaggs et al., 2014). In fish brain, the GFAP+ cell processes are more likely to restore axons than to form any glial scar (Goldshmit et al., 2012; Takeda et al., 2015).

### 4.3. Neuronogenesis in the cerebellum of juvenile O. masou after traumatic injury

The investigation of distribution of HuCD+ cells in the cerebellum of juvenile *O. masou* revealed a significant increase in the number of immunopositive cells in the dorsal part of cerebellum after injury. These data are consistent with those obtained from other fish species (Kaslin et al., 2009; März et al., 2011). We believe that the increase in the number of new neurons in DMZ of CCb in juvenile *O. masou* is a reaction to injury; a similar functional response was observed in juvenile *O. masou* after a telencephalon injury (Pushchina et al., 2017a), as well as in adult trout after an optical nerve injury (Pushchina et al., 2016).

In the post-injury period, a large number of HuCD+ cells were detected throughout the dorsal zone. In the control fish, there were rounded type I cells forming constitutive neuroepithelial neurogenic niches in the surface of lateral and basal zones; after the injury, HuCD+ cells were detected in the reactive neurogenic domains containing heterogeneous cell populations of all types, including the large, intensely labeled, elongated cells. The increase in the number of reactive neurogenic domains containing HuCD+ cells, in the intensity of HuCD labeling of such cells in the neurogenic zones of the surface part of CCb molecular layer, and in the number of radially migrating cells was significant (Pushchina et al., 2018). After the damage to the pallial zone of the telencephalon in juvenile *O. masou*, we also observed an uneven increase in the number of HuCD+ cells in the dorsal, middle, and lateral zones accordingly (Pushchina et al., 2017a). Thus, the highest number of HuCD+ cells was found in the lateral part of the dorsal pallial zone, where we also observed the most intensive cell migration (Pushchina et al., 2017a). After the cerebellar injury, an increase in the number of HuCD+ cells was also detected in the basal part of cerebellum compared with the control animals. However, in the basal part, along with small undifferentiated HuCD+ cells, the number of differentiated HuCD+ cells with processes and ganglionic cells (Purkinje cells and EDC) was also found to increase. In the post-injury period, a large number of HuCD+ cells were distributed throughout the dorsal zone. In the control animals, at the surface of the lateral and basal zones, there were rounded (type I) cells forming constitutive neuroepithelial niches; after the injury, HuCD+ cells were detected in the reactive neurogenic domains containing heterogeneous cell populations of all types, including large, intensely labeled, elongated cells. Thus, it is possible that a traumatic damage to cerebellum not only enhances neuronogenesis in reactive neurogenic niches, but also may accelerate neuronal differentiation of cells, which ingresses the mature neuronal networks of the cerebellum. A retrograde tracing in combination with BrdU labeling of cells showed that the processes of the new granular neurons of zebrafish are projected into the molecular layer (Zupanc and Ott, 1999). This fact allows us to assume that these neurons are integrated into the already existing neural network in cerebellum.

One of the first responses to a CNS trauma in *D. rerio* is the activation of immune cells (Finsen and Owens, 2011; Ransohoff and Brown, 2012) and radial glia, a significant increase in proliferation, and the creation of new neurons to replace lost ones (Kroehne et al., 2011; Kizil et al., 2012). A short-term labeling of BrdU and/or PCNA labeling revealed extensive proliferation in the molecular layer of zebrafish cerebellum (Grandel and Brandt, 2013). The migrating progenitors and the differentiating granular neurons express different transcription factors: Zic1, Zic3, and Pax3 (Kaslin et al., 2009).

Thus, after injury, the newly formed cells in the cerebellum of juvenile *O. masou* are formed from two sources of endogenous stem cells located near the injury zone. The first source is the dorsal proliferative zone, which contains neurons formed during constitutive neurogenesis. The level of cell proliferation in DMZ slightly increases after damage, as compared to the intact animals. The second source is an area where proliferative activity occurs only after injury; in the intact brain, it is mitotically “silent” (Pushchina et al., 2018). Such zones correspond to reactive neurogenic niches arising after trauma in various parts of the molecular layer, mostly in the caudal part of cerebellum. Within a few weeks after injury, neurons are eliminated by apoptosis and replaced by new cells (Zupanc, 1999). Within the first ten days, the rate of cell proliferation in the damaged site increases multifold compared to other parts of the cerebellum body. The diffuse pattern of cell death is a characteristic feature of cerebellum in the post-traumatic period in juvenile *O. masou* (Stukaneva et al., 2015; Pushchina et al., 2017b). A damage to the blood-brain barrier is accompanied by an intense inflammatory response, both from the side of microglia and from monocytes, macrophages, T-cells, and neutrophils invading the injury area (Raivich et al., 1999).

### 4.4. Glutamine synthetase in the cerebellum of juvenile O. masou after injury

Glutamate is a major exciting neurotransmitter in the CNS (approximately 8–10 mM/kg) and is found in approximately 80% of all neurons (Revett et al., 2013). It participates in most normal brain functions and plays an important role in the development of CNS, in the elimination of synapses, cell migration, differentiation, and death (Komuro and Rakic, 1993). Most of glutamate in vertebrate brain is localized intracellularly, as well as in nerve fibers and endings. Only a small part of it is present in the intercellular space (Gasparini and Griffiths, 2013). In mammals, neurotransmitters of cerebellar neurons have been well investigated. Purkinje cells, stellate cells, basket cells, and Golgi cells are inhibitory neurons that mainly use GABA as a neural transmitter, whereas granule cells are excitatory neurons that use glutamate (Ottersen, 1993).

Removal of glutamate from the intercellular space is the main cause of changes in glutamate-mediated neurotransmission, receptor activation, induction, and depolarization of neurons, whose activity is regulated by the intracellular level of Ca^2+^ and Na^2+^ fluxes (Revett et al., 2013). These processes lead to exocytosis of glutamate and immediate cell death, which correlates with the impairment of memory functions and training in case of disorders of these processes (Wenk et al., 2006).

In the brain of adult zebrafish, GS is not expressed in the proliferative zones of optic tectum and dorsolateral telencephalon (Schmidt et al., 2013). In these areas, glial cells are stained with BLBP, a marker of early differentiation of glial cells (Campbell and Götz, 2002; Raymond et al., 2006). The lack of GS in the embryonic zones is probably explained by the absence of glutamate excretion in them, and, as a result, the GS is not detected at an early stage of differentiation of astroglial cells. However, this indicates a functionally corrected expression of the protein in these cells (Grupp et al., 2010). NSCs in the constitutive neurogenic niches of the ventricular regions are devoid of radial processes and, as supposed, create neurons and glia. In contrast, proliferating PCNA+ cells in the ventricular zone differ from proliferating cells in the embryonic zones and are GS-immunopositive (Grupp et al., 2010). These cells have long processes and probably correspond to RG cells capable of creating neurons and glia during the development of mammalian CNS (Campbell and Götz, 2002).

The analysis of IHC activity of GS in the control animals and after a damage to cerebellum of juvenile *O. masou* revealed significant differences in distribution of the enzyme. In both cases, GS activity was identified both in cells and in fibers. The results of the morphological analysis and recent data (Revett et al., 2013) indicate that GS-containing cells are populations of astrocyte-like cells. The morphological studies conducted on *O. masou* juveniles show the presence of a heterogeneous population of GS+ cells in the control. In the intact animals, in the area of molecular layer of *O. masou* cerebellum, a GS activity was detected in type VI and VII cells. The density of distribution of GS+ cells in the molecular layer in the intact animals was quite high, which indicates a high level of enzyme activity and, possibly, its influence on the processes of constitutive neurogenesis.

After the injury, a 1.5-fold increase in the number of intensely labeled, large type VII cells, compared to the control, was recorded. The number of intensely GS+ labeled type VII cells in the dorsal, lateral, and basal zones after the injury increased 1.9-fold. Type VI cells with moderate GS activity were located singly, and more intensely labeled type VI cells formed clusters. The appearance of new GS+ cells and their aggregations after injury was more characteristic for the molecular layer of CCb.

After the injury, in all the areas of CCb, the amount of GS+ fibers increased. In this case, the activity of fibers in the granular layer (*eminentia granularis*) was normal, not high. Such a complex redistribution pattern of GS activity after the injury indicates significant reorganization of enzymatic activity, both in fibers and in cells.

Thus, after the injury, in the molecular layer of CCb of juvenile *O. masou*, we observed a change in the cell composition of GS+ elements and a general increase in the number of GS+ cells of various types (1.6-fold) and GS+ fibers (1.2-fold) on day 3, compared to the control. The number of GS+ cells gradually increased on day 3 after the traumatic injury.

We detected the presence of GS-positive elements, including both heterogeneous populations of cells and fibers. We believe that GS in the cerebellum of juvenile *O. masou* can label both glutamatergic neurons and astrocyte-like cells, containing GS and detoxifying glutamate re-uptake from the intercellular space (Pushchina et al., 2017b). It was found that after a treatment with methionine, the activity of GS in the brain of zebrafish does not change. A decrease in glutamate uptake was probably not associated with a decrease in its amount, and treatment with methionine did not lead to changes in glutamatergic neurotransmission in zebrafish brain (Vuaden et al., 2016).

During a morphological observation of the cerebellum of juvenile *O. masou*, we also identified GS-containing fibers. In the basal part of cerebellum, particularly large and thick fibers with a high level of GS activity were detected. Such fibers often contained large GS-immunopositive endings with a high level of enzyme activity. Based on the distribution of ascending afferents in the cerebellum of fish (Wullimann, 1998), we assume that this type of fiber can correspond to the ascending glutamatergic liana-like fibers of other vertebrates, whereas the identified end thickenings are the ending fibers of nodular type. This assumption is also confirmed by the identification of glomerula-like complexes in the granular layer of cerebellum. According to our results, formations containing a large number of GS-immunopositive presynaptic terminals are present in the granular layer of CCb in juvenile *O. masou* (Pushchina et al., 2017b). Taking into account that no analogous complexes containing other mediator systems have been known to date, we are inclined to attribute these complexes, exhibiting the GS activity in the granular layer of CCb in juvenile *O. masou*, to glomerular-like formations similar to the mammalian cerebellar glomeruli.

Concerning the synaptogenesis of the granular layer, it appears that, as in other vertebrates, the mossy fibers, the granule cells, and the Golgi cells participate in the formation of glomeruli, which are the synaptic complexes of this layer (Pouwels, 1978c). Mossy fibers contain more neurofilaments than other axons in the cerebellum of trout and can easily be recognized. In trout, 17-mm mossy fiber synapses are observed for the first time. In this stage, the rosette has a rather simple form (Pouwels, 1978c) and contacts with only a few dendrites of granule cells. These endings contain elliptical vesicles. In accordance with light microscopic observations and with data on the structure of the mammalian glomerulus available in literature, these terminals are referred to as axons of Golgi cells. A glomerulus is surrounded by glial processes (Pouwels, 1978d).

We believe that among heterogeneous population of GS+ cells in juvenile *O. masou* there are both glutamatergic neurons containing GS as an enzyme-metabolizing glutamate and astrocyte-like cells that can receive glutamate as a result of its re-uptake from the intercellular space. Observations show a significant change in the GS activity in various parts of the cerebellum after a damaging impact on it in juvenile *O. masou*. Such spatial specificity can be associated, on the one hand, with both the toxicity effects caused by the injury and the change in glutamatergic neurotransmission in the damaged neuronal networks. On the other hand, the high activity of GS in the cerebellum of intact juvenile *O. masou* clearly indicates the participation of glutamate in processes of neuronal plasticity, particularly in the morphogenesis that takes place during the constitutive neurogenesis.

We suggest that metabolic glutamate, which is presumably involved in the morphogenetic functions of the cerebellum in juvenile *O. masou*, under normal conditions can be localized both in growing neurons and in fibers (Pushchina et al., 2017b). This assumption is supported by the results of the studies, which confirm the high activity of GS in cells of the molecular layer, especially in the dorsal part of CCb. This area of *O. masou* cerebellum contains DMZ, which is distinguished by high neurogenic activity in the control animals. After the injury, the GS activity increases; however, we do not rule out that these changes are not generalized, but are a particular display of the change in GS activity on day 3 post-injury. This hypothesis is also confirmed by the data of the enzyme immunoassay (ELISA), carried out by us on cerebellum of juvenile *O. masou*. Thus, in the course of long-term monitoring, a gradual increase in GS activity was observed on day 2 and its slight decrease on day 3 after the cerebellar damage (**Figure 10**). Nevertheless, these changes in the metabolic activity of GS were only a local decrease in the activity of the enzyme, displayed on day 3. As a result of ELISA, it was found that the activity of the enzyme after the damage manifests a complex, multi-peaked pattern. An increase in the activity of the enzyme was observed on days 1, 2, and 5 post-injury. On day 3, according to ELISA data, there was a slight decrease in the GS activity. Thus, a decrease in the GS activity on day 3 after the damage to the cerebellum of juvenile *O. masou* may be a particular display of a change in the metabolic status. Due to a sufficiently high intensity of constitutive neurogenesis observed in juvenile *O. masou*, we tend to believe that the response from GS-producing elements in the cerebellum can be complex. Thus, a damaging effect can cause the death of a large number of cells in the cerebellum of *O. masou* by apoptosis (Stukaneva et al, 2015; Pushchina et al., 2018), due to glutamate toxicity. However, the increased production of GS on days 1, 2, and 5 indicates a sufficiently high level of production of the enzyme, whose activity can decrease cyclically and be determined by various factors, the nature of which is still unknown.

The increase in the number of GS+ cells in the molecular and granular layers and the increase in the activity of the enzyme on day 3, observed by us, can be attributed to the astrocyte response observed during the damage. However, the concentration of such cells is not high enough to relate this response of GS+ cells to the formation of an astrocytic response like that after an injury in mammalian CNS. As is known, a pool of reactive astrocytes, with their morphological and biochemical features being significantly different from normal astrocytes, is formed after a traumatic impact on brain of mammals (Revett et al., 2013). The cellular mechanisms associated with the transformation of astrocytes population and the allocation of a subpopulation of activated astroglia in brain of fish have not been studied yet. In contrast to astrocytes of mammals, those in fish brain do not form the so-called astrocytic barrier (Baumgart et al., 2012; Goldshmit et al., 2012; Takeda et al., 2015), which is characteristic for the development of the posttraumatic process in the mammalian brain (Silver and Miller, 2004; Sofroniew, 2009). Nevertheless, the change in the synthesis of GS clearly confirms the neuroprotective properties of this enzyme and its increased production in the cerebellum of juvenile *O. masou*. Thus, GS is not only a marker of cells involved in the conversion of glutamate/glutamine, but it can also be considered an effective neuroprotection factor that promotes and facilitates the post-traumatic reparative process.

According to Zupanc and Sîrbulescu (2013), in the cerebellum of *Apteronotus leptorhynchus* after injury, the amount of GS increases, while the synthesis of this enzyme in CNS of mammals after damage is suppressed (Grosche et al., 1995). The success of regenerative processes in the fish brain after injury is determined by a number of factors that distinguish the dynamics of this process from those in mammals. According to Grosche et al. (1995), GS is a specific glial protein that converts toxic glutamate, accumulated as a result of neuronal damage, to a non-toxic amino acid, glutamine. Under normal conditions, this mechanism prevents the neurotoxic accumulation of glutamate in the neural tissue, protecting the neurons from cell death. But after a brain injury in mammals, the amount of synthesized GS is insufficient to neutralize the toxic effects of glutamate. This has such consequences as the development of primary and secondary inflammation, as well as progressive neurodegenerative processes observed after a CNS trauma in mammalian animals and humans. In fish, however, the enhancement of GS activity probably provides an important mechanism for reducing the neurodegenerative effects caused by the glutamate neurotoxicity. This assumption is based on the presence of certain mechanisms that determine significant differences in regenerative abilities between mammals and fish (Zupanc and Sîrbulescu, 2013).

In mammals, the reactive gliosis, trauma-induced inflammation, and glial scar formation are considered the main obstacles to successful brain regeneration (Silver and Miller, 2004; Fitch and Silver, 2008; Sofroniew, 2009). One of the therapeutic approaches, developed to protect cells from degeneration, is based on increasing the GS in glial cells by inducing the expression of an endogenous gene or an exogenous stock of purified enzyme in a mammalian retina explant culture (Gorovits et al., 1997).

In the telencephalon of zebrafish (Ganz et al., 2010; Grupp et al., 2010; März et al., 2011), as well as in *Cyprinus carpio* brain (Kálmán, 1998) and *Chelon labrosus* brain (Arochena et al., 2004), no stellate astrocytes typical of mammals were found. In the telencephalon of adult *D. rerio*, the radial glia, like the mammalian astroglia, exhibits characteristic signs of reactive gliosis immediately after injury (Vitalo et al., 2016). Despite reactive astrocytes do not exhibit neurogenic properties in the brain of adult mammals, they can give rise to multipotent neurospheres and neurons *in vitro*, which emphasize the important role of a restrictive, inadmissible non-permissive medium *in vivo* (Costa et al., 2010; Heinrich et al., 2010; Robel et al., 2011). In contrast, the reactive radial glia of zebrafish performs functions of endogenous neurogenic progenitor cells after damage. The radial glia creates neuroblasts that migrate both into the constitutive, periventricular region and into the deep layers of parenchyma to the site of injury, which is usually not observed in intact animals (Vitalo et al., 2016).

